# D2-expressing neurons of the anterior paraventricular nucleus of the thalamus as key players for modulating relative aversive and safety learning

**DOI:** 10.1101/2024.12.30.630766

**Authors:** Magdalena Miranda, Elsa Karam, Pola Tuduri, Maïwenn Passard, Majda Drahmani, Loic Vaillant, Christina-Anna Vallianatou, Marina Lavigne, Eric J. Kremer, Jeanne Ster, Emmanuel Perisse, Stéphanie Trouche

## Abstract

Associative learning provides a means to assign appropriate value signals to experiences that animals can later use to make appropriate value-based choices. By generalizing from prior aversive experiences and comparing with a current experience, animals can learn the relative aversive (“better or worse than”) value or relative positive safety value between experiences. Yet, how such relative value signals are acquired during learning remain poorly known. The paraventricular nucleus of the thalamus (PVT) is critical for the control of salience and valence signals. It is therefore well-placed to be involved in learning relative aversive value and learned safety. Using a newly-developed conditioned place preference task, we uncover specific behavioral signatures associated with learning relative aversive value and learned safety, and related value-based choices. We then show that neurons expressing dopamine D2 receptor (D2+) in the anterior PVT (aPVT) are preferentially recruited by both relative aversive value learning and learned safety. Using a pharmacogenetic strategy, we demonstrate that aPVT D2+, but not D2-, neurons are involved in learning relative- but not absolute- aversive value. Finally, we show that aPVT D2+, but not D2-, neurons are also critical for the encoding of learned safety. Overall, our findings reveal a novel role of aPVT D2+ neurons in computing relative aversive/safety value signals between experiences during learning to promote appropriate value-based choices.

## Introduction

Learning that some cues can predict threat while others can predict safety is critical for appropriate decision-making and survival. ^1–3^ By generalizing predictions ^4,5^ and comparing the value of contextual/sensory signals of a prior aversive experience with an ongoing threat, animals can learn the relative aversive value between two experiences (i.e. whether one is better or worse than the other). ^6–11^ However, by generalizing prediction and comparing a prior aversive experience with the absence of a threat, animals can learn relative safety. ^12–14^ Moreover, previous studies showed that learning the relative aversive value between two aversive experiences, such that one is perceived as better than the other, involves the reward system.^6,7,15^ Similarly, a safe experience, relative to a prior aversive one, can acquire a relative positive value called learned safety, a process that also requires the reward system. ^16,17^ When later confronted with a choice between two options, pre-computed values acquired during learning (here relative aversive value or learned safety) facilitate decision-making processes, thus promoting more efficient and accurate value-based choices. ^18^ Critically, overgeneralization and dysregulation of a normal threat and safety learning, leading to alteration of relative value computation, is maladaptive and serves as a major feature of fear-related disorders including post-traumatic stress disorder (PTSD). ^4^ While fear generalization ^4,19^ and the interaction between appetitive and aversive systems ^7,20–22^ are likely crucial players in the encoding of value signals, how the comparison of generalized predicted and current values contribute to the encoding of relative aversive value and learned safety remains poorly understood.

The paraventricular nucleus of the thalamus (PVT) is a hub connecting the thalamus with cortical and limbic structures ^23–27^, which is critically involved in arousal, stimulus salience processing and value assignment to associative stimuli. ^28,29^ ^30–33^ The PVT is therefore a strong candidate structure for disentangling motivational conflicts and comparing experiences to compute appropriate relative value signals during learning. ^24,28,33–37^ Due to its involvement in granting and updating the value of aversive experiences ^28,36,38–42^, here we hypothesize that the PVT is crucial for the assignment of a relative aversive/safety value. Of note, the D2 dopamine receptor-expressing neurons (D2+) of the PVT are sensitive to valence ^33^ and are implicated in signaling aversive states. ^33,39^ By contrast, D2-non-expressing neurons (D2-) are modulated by arousal, relaying a salience signal to the cortex. ^33^ According to this functional dissociation, D2+ neurons could be critical to signal a “better than expected” relative aversive value. The PVT is also instrumental for the correct assignment of a learned safety value to an experience. ^42^ However, it remains unclear whether D2+ neurons of the PVT differentially contribute to encoding relative aversive value and/or learned safety.

Here, we addressed this question by designing a novel conditioned place preference task that allows mice to encode either relative aversive value or learned safety. In this task, mice learn to associate one context with foot shock delivery, followed by a second context that is either associated with shocks of lower intensity (learning a relative “better” aversive value), or with the absence of shocks (for learned safety). After learning, mice are confronted to a choice between the two contexts. We showed that mice adapt their defensive responses to each task, exhibiting distinct behavioral signatures during learning and choice tests. Our data revealed that during learning of the second context, when comparing it to the previous aversive experience, both tasks preferentially recruit D2+ neurons in the aPVT compared to pPVT. We then found that pharmacogenetic inactivation of aPVT D2+, but not D2-, neurons during the exposure to the second context disrupt relative aversive comparison by increasing its aversiveness. This led to a suppressed preference towards the less punished-associated context during the choice test. Consistently, when mice learn to associate two distinct contexts paired with the same shock intensity, inactivation of aPVT D2+ neurons during the exposure to the såecond context also increases its aversiveness, tilting mouse preference towards the first context during choice test. Finally, we found that inactivating aPVT D2+ neurons increases safety of the ongoing experience inducing enhanced preference towards the safe context during choice test.

Taken together, our data demonstrate that aPVT D2+ neurons modulate the comparison of values during learning that enables accurate assignment of relative aversive value and learned safety, thereby promoting appropriate value-based choices. These results also provide compelling evidence that aPVT neuronal ensembles expressing D2 receptors, embedded within limbic-cortical circuits, are key in the evaluation process for assigning correct value signals to sensory experience during associative learning.

## Results

### Mice use different behavioral strategies for learned relative safety and relative aversive value-based learning and choice

To study the neural circuits and mechanisms underlying learned safety and relative aversive value coding, we designed a novel 1-day conditioned place preference (CPP) task **(Figures 1A** and **S1A**). In this task, mice are subjected to a pretest session ("Pretest"), allowing them to explore two never-encountered contexts (A and B connected via a bridge) followed by two conditioning sessions ("S1" and "S2") in each context, and a test session ("Choice test") between contexts A and B. Three groups were tested. In the control group, no shock was delivered in any context during S1 and S2. To study learned relative safety value (named the “safe” group), we employed a discriminative/differential fear conditioning paradigm ^12–14^ during which mice received four electric shocks of 0.6 mA in context A during S1 ("A+0.6"), but no shock (no threat) in context B during S2 ("B+0"). To study learned relative aversive value (named the “aversive” group), mice were subjected to four electric shocks of 0.6 mA in context A during S1 ("A+0.6") as for the safe group, but received four electric shocks of 0.3 mA in context B during S2 ("B+0.3"). As expected, no freezing was observed during Pretest in any group (**Figure S1B**). Both safe and aversive groups exhibited increased freezing levels in S1 and S2 conditioning sessions compared to the control group (**Figure S1C**). However, only mice in the aversive group showed increased freezing from S1 to S2 **(Figure 1B**), along with heightened anxiety as measured with a decreased time spent in the center during S2 (**Figure S1D**). By analyzing the dynamic of freezing during S1 and S2 (**Figures S1E**), we found that freezing before the putative first shock of S2, but not S1, significantly increased for both safe and aversive groups compared to control group (**Figure 1C**). The increased freezing observed at the beginning of S2 suggests a generalized expectation based on their previous aversive experience in context A+0.6 mA. Importantly, while both aversive and safe groups increased freezing after the 4th (and last) shock in S1 (**Figures 1D, S1E** and **S1F**), only mice in the aversive group maintained a high freezing level at the end of S2 (i.e., after receiving four shocks in context B), indicating that the aversive group learned that context B is aversive (**Figures 1D** and **S1E**). In contrast, mice in the safe group significantly decreased their freezing after the fourth (and last) putative shock (**Figure 1D**) suggesting that mice start to discriminate both contexts, perceiving context B+0 mA as relatively safer than context A + 0.6 mA. Strikingly, we also observed tail rattling behavior only in the aversive group immediately after each shock event in S2 (**Figure 1E; video 1**), but not S1 (**Figure S1G**). Thus, the increased number of shock events shifted mouse’s defensive responses. The absence of such active defensive behavior in response to a threat ^43^ in the safe group strengthen the idea that context B + 0 mA became relatively safer than context A+0.6 mA. Overall, our results indicate that relative to a prior intense aversive experience, mice assign a relative aversive/safe value to ensuing less punished/not punished experience.

**Figure 1.**
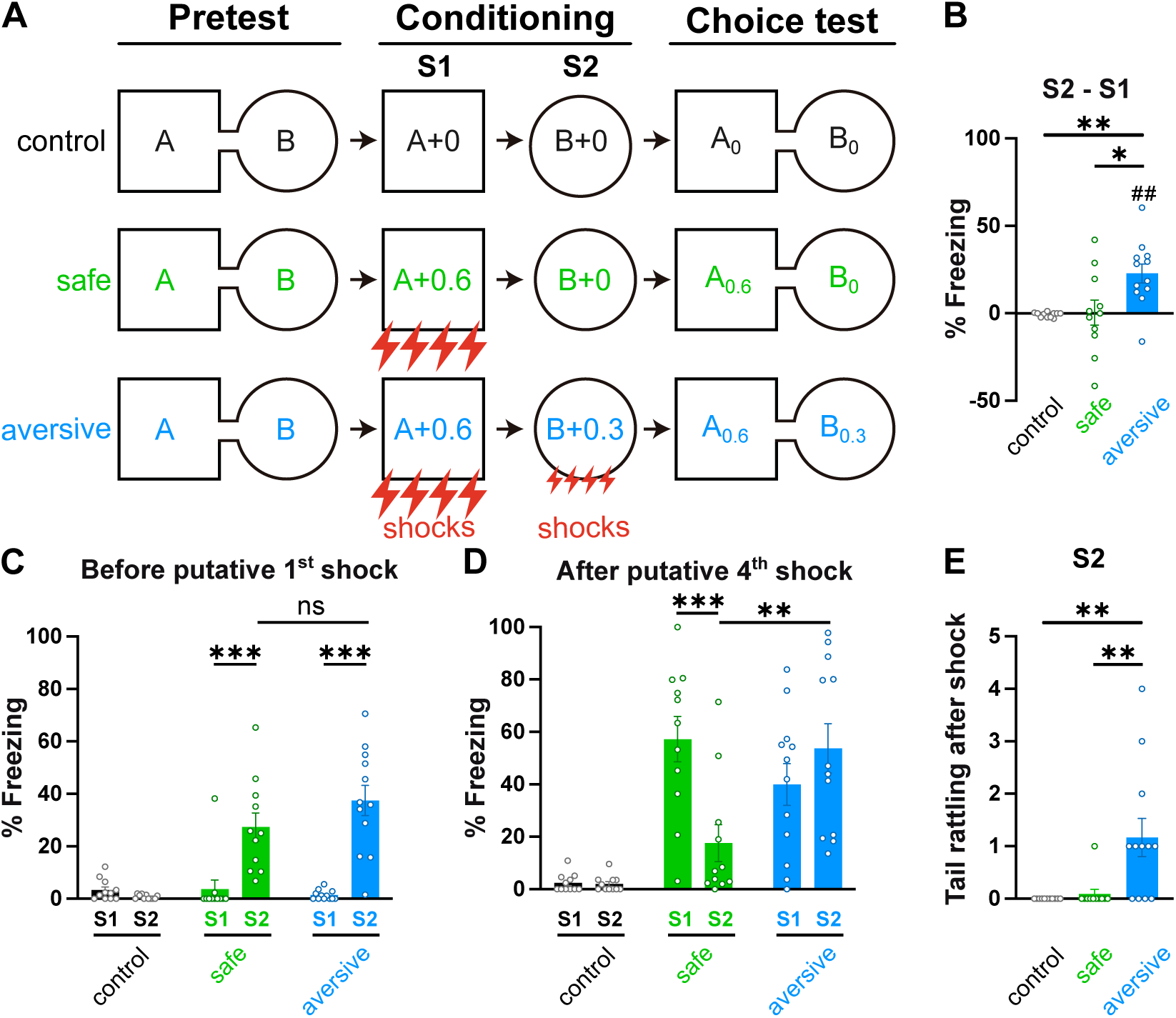

One hour after the learning phase, all mice were tested during a choice-test session. Importantly, mice were first placed in their most preferred context (defined during Pretest), i.e., context A0 for the control group and the most punished Context A0.6 for both safe and aversive groups (**Figure 2A)**. Their latency to cross from context A to context B (i.e., First latency) was similar between all 3 groups during choice test (**Figure S2A**). Similarly, the number of tail rattling was similar before their first cross to context B (**Figure S2B; video 2**). However, only mice from the aversive group spent more time in the bridge before the first cross (**Figures 2B** and **2C**). In addition, the majority of mice in the aversive group (10 out of 12) displayed a "turn-back" behavior in the bridge zone (**Figure 2D; video 3**). Such behavioral signature is reminiscent of the “vicarious trial and error” behavior with animals moving back and forth in direction of different choices at decision zones. ^44,45^

**Figure 2.**
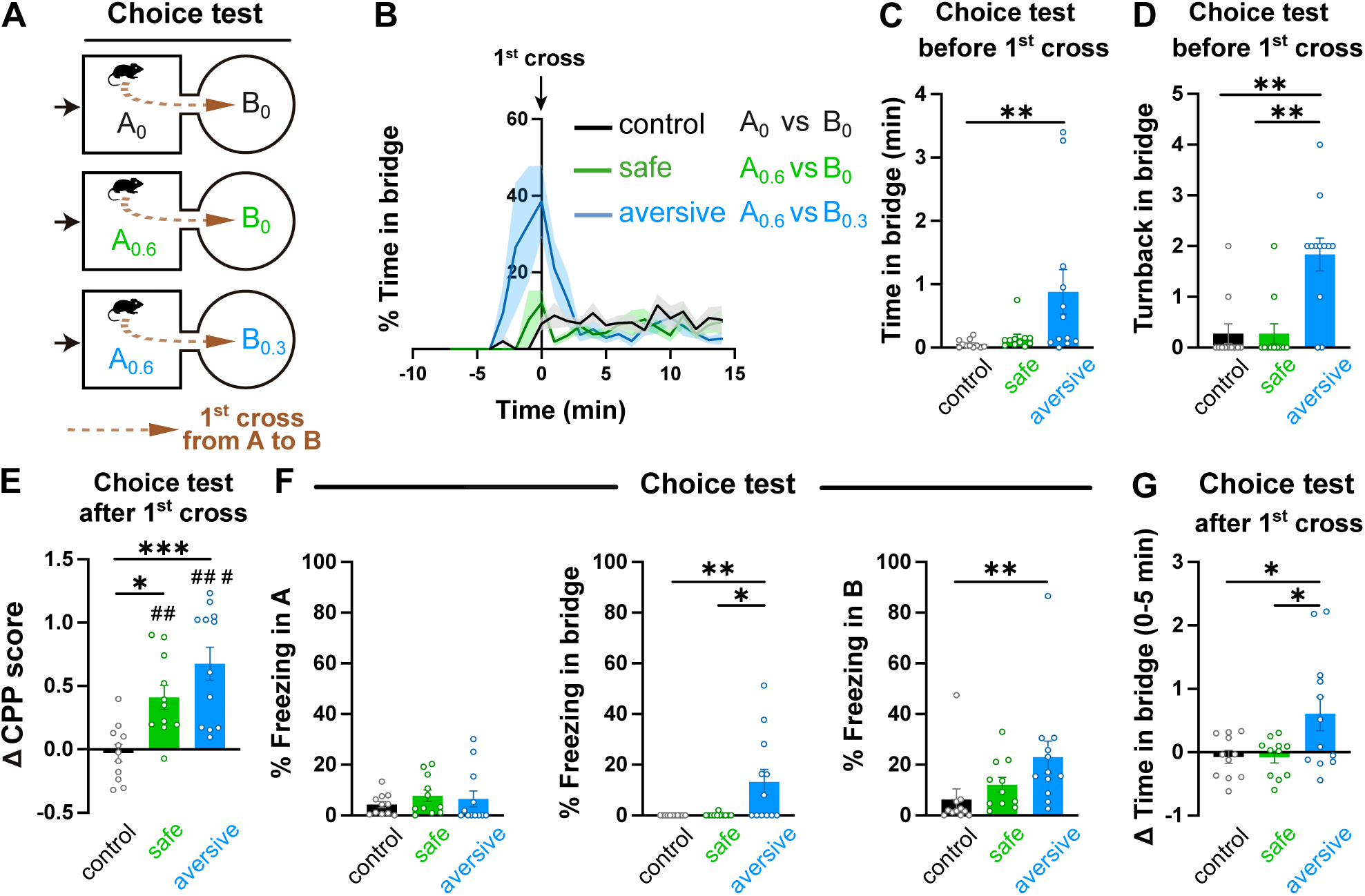

After crossing the bridge for the first time (i.e., First cross), mice were allowed to explore the two contexts for 15 min (as in ^46,47^) (**Figure S2C**). We calculated a ΔCPP score comparing the time spent in each context during pretest (15 min, **Figure S2D**) vs choice test (15 min after first cross, **Figure 2E**). As expected, control mice showed no preference for either of the two contexts during choice test (ΔCPP score close to zero, **Figure 2E**). Mice from the safe and aversive groups similarly reverted their initial preference and spent more time in the less aversive context (positive ΔCPP score significantly different from zero and not significant between safe and aversive groups, **Figure 2E**), along with a reduced exploratory behavior (**Figures S2C** and **S2E**). Their preference was similar for both safe and aversive groups during the first 5 min of the test, but was sustained in the aversive group throughout the choice test (**Figure S2F**). For the aversive group, this preference was accompanied with increased freezing in the bridge zone and context B0.3 (**Figure 2F**), along with higher stress marked by more feces (**Figure S2G**). In line with these results, the aversive group stayed longer in the bridge after the first cross (**Figure 2G**). These data suggest that the aversive group therefore utilized prior learning information to avoid the most punished context A0.6 and potentially approach context B0.3 ^15^ while being anxious and showing hesitation to make a choice between the two aversive contexts. By contrast, the safe group preferred the safe B0 context and showed less anxiety-like behavior. Overall, mice use learned experience to adopt specific behavioral signatures and enable appropriate value-based choices.

### Both learned relative safety and relative aversive learning preferentially recruit D2+ neurons in the aPVT

We hypothesized that mice compare the relative value of the two experiences during conditioning S2 relative to S1 and assign either a relative “better than” aversive value to context B (paired 0.3 mA shocks) for the aversive group ^6,7,34^ or a relative safety value to context B (no threat) for the safe group. ^12,14,17^ However, it remains unclear how mice encode a relative value (aversive or safe) between experiences, and how this influences subsequent value-based choices. The PVT is central for modulating anxiety-related conflicting appetitive-aversive information ^37,48^ and could be pivotal for the encoding of relative (aversive or safe) value. Notably, the functional PVT antero-posterior axis expressing dopaminergic D2 receptor (D2+ neurons) involved in salience and valence coding ^33,49,50^ could extricate the different intensities of prior (during S1) and actual information learned during the conditioning session S2. We first tested this hypothesis by asking whether PVT neurons, in particular D2+ neurons, are differentially recruited during learning using D2 Cre:Ai14 mice (see Methods). First, our *ex vivo* whole-cell recording experiments confirmed that td-Tomato (i.e., Ai14) cells respond to quinpirole (D2R-like agonist, 1□M; action potential (AP) frequency baseline = 1.00 +/- 0.20 Hz, AP frequency quinpirole = 0.37 +/- 0.21 Hz, n= 5 cells, paired t-test p= 0.0047 **) validating the use of this mouse line. We then performed a cFos immunostaining in the aPVT and pPVT in D2-tagged animals of 3 groups: home-cage (HC), safe and aversive (**Figures 3A** and **3B**). To reflect cFos activity at the time of the comparison between the two conditioning sessions, mice were perfused 90 min after the conditioning session S2. Importantly, we verified that both safe and aversive groups showed distinct freezing patterns at the end of S2 (**Figures S3A** and **S3B**), as we previously described (**Figures 1C** and **1D**). We found that both safe and aversive groups show similar, and significantly higher, levels of activated (cFos positive) neurons compared to the home-cage group (**Figure 3C**). In the safe and aversive groups, such recruitment was more prominent in the aPVT than the pPVT (**Figures 3C** and **S3C**) and incorporated a higher proportion of D2+ than D2- neurons in the aPVT, but not pPVT (**Figures 3D**-**3E**). Noteworthy, D2+ cell densities were similar between all groups, but were higher in pPVT than aPVT **(Figure S3D)**, as previously published. ^33^ These results indicate that both learned safety and relative aversive learning recruit preferentially aPVT D2+ neurons. Mice were subjected to a choice test immediately before being perfused (**Figures 3A**). In line with our previous findings (**Figure 2E)**, both safe and aversive groups significantly preferred the non-punished (context B0) or less punished compartment (context B0.3) during choice test, respectively (positive ΔCPP score in **Figure S3E**). As shown in **Figure 2**, the relative choice between the two previously (and differently) punished contexts, A0.6 and B0.3, led mice to spend a prolonged period in the bridge, likely assessing the two aversive options to make an appropriate choice (**Figure S3F**). Given the role of the PVT and the increased activation of D2+ neurons during the conditioning session S2, we hypothesized that mouse behavior during the choice test is closely linked to the activity of aPVT D2+ neurons during S2. Accordingly, the percentage of activated D2+ neurons of the aPVT (but not pPVT) of the aversive group during S2 is negatively correlated with the time spent on the bridge (i.e., the decision zone) during the choice test (**Figure 3F**). In other words, the less aPVT D2+ neuron activation during S2, the more time mice will spend in the bridge at the start of the choice test. Strikingly, such correlation was not observed for aPVT D2-negative activated neurons (**Figure 3F**), nor for the safe group (**Figure 3G**). These results strongly suggest a crucial role of aPVT D2+ neurons for learning relative aversive value during conditioning to subsequently promote an appropriate relative aversive value-based choice.

**Figure 3.**
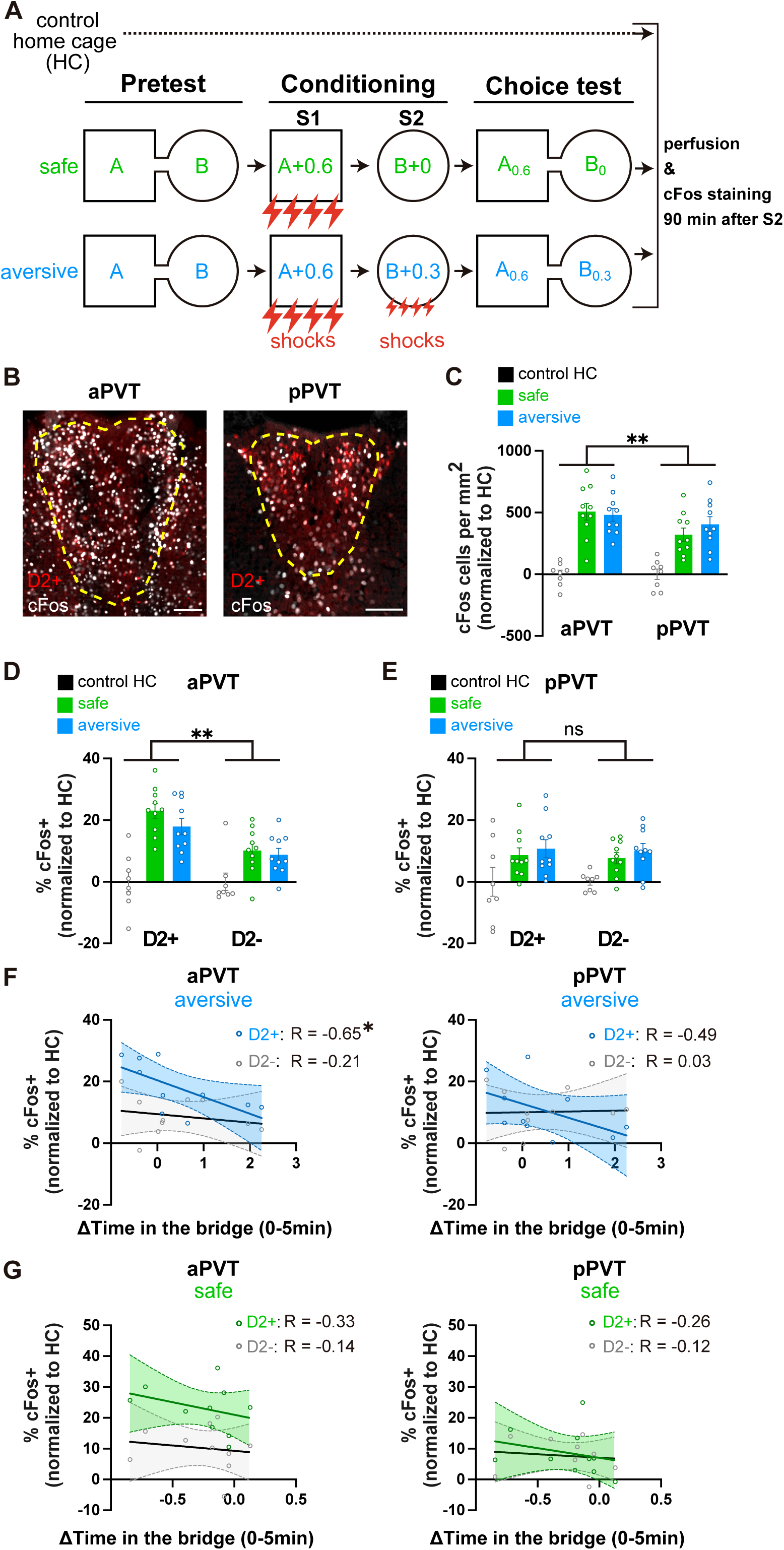

### aPVT D2+, but not D2-, neurons are crucial for learning relative aversive value

To directly test the role of aPVT D2+ neurons for learning relative aversive value during conditioning in S2, we virally injected D2 Cre mice with viral vectors expressing a Cre-dependent DREADD (AAV-DIO- hM4Di-mCherry) or control viral vector (AAV-DIO-mCherry) in the aPVT (**Figure 4A**). Four weeks later, mice underwent the aversive CPP task and were injected with Clozapine N-Oxide (CNO) intraperitoneally 30 min before S2. This pharmacogenetic strategy aimed to inactivate aPVT D2+ neurons when mice relatively compare the S1 and S2 experiences (**Figure 4B**). To ascertain the effect of hM4Di activity, we performed *ex vivo* whole-cell recordings in control- and hM4Di-D2+ neurons of the aPVT, identified by their mCherry fluorescence (**Figure S4A)**. The application of 10□M CNO to aPVT brain sections expressing inhibitory DREADD (hM4Di) hyperpolarizes D2+ neurons by drastically decreasing spike frequency (**Figure S4A)**. By contrast, CNO had no impact on aPVT D2+ neurons transduced with the control AAV-DIO-mCherry virus (**Figure S4A)**. In line with previous reports, we also confirmed projection targets of aPVT D2+ neurons such as nucleus accumbens (NAc) and amygdala (**Figure S4B**). ^33^ First, and as expected, both control and hM4Di groups showed a similar preference of context during pretest before CNO injection (**Figure S4C).** While silencing aPVT D2+ neurons during S2 did not change the amount of freezing during conditioning (**Figures S4D** and **S4E**), it led to a significant increase of tail rattling behavior in S2 for the hM4Di group immediately after receiving electric shocks (**Figure 4C**). Moreover, while we observed a decreased average distance travelled after shock delivery between S1 and S2 in the control group (**Figure 4D**), inhibiting aPVT D2+ neurons prevented such decrease during S2 (**Figure 4D**). These findings suggest that silencing aPVT D2+ neurons heightened shock perception during S2, thereby possibly increasing the aversiveness of context B+0.3. As a consequence, before their first cross from context A0.6 to context B0.3 during choice test, mice displayed a higher number of turnbacks (**Figure 4E**) and tail rattling (**Figure 4F**) in the bridge without changing their first latency to cross (**Figure S4F**) nor the time they spent in the bridge zone (**Figure S4G**). These results indicate that hM4Di mice perceived context B0.3 as threatening as context A0.6. In line with this conclusion, the preference of control mice for the previously less punished context B0.3 was fully suppressed in hM4Di mice (ΔCPP score close to 0; **Figure 4G**). This was true when aPVT D2+ neurons were inactivated during S2, but not during choice test, highlighting the selective role of aPVT D2+ neurons for relative aversive value-based learning (**Figure S4H-J**). hM4Di mice spent more time in the bridge zone after their first cross from context A0.6 to context B0.3 (**Figure 4H**), indicating more hesitation to choose between the two contexts. Moreover, mice froze less in context B0.3 (**Figure 4I**; also see **Figure 2E**). Overall, our results converge towards the idea that silencing aPVT D2+ neurons enhanced the aversiveness of the “context B+0.3 mA” learning experience to reach a similar aversiveness as in context A+0.6 mA. This would therefore prevent mice from accurately learning the relative difference between two aversive experiences and from assigning a relative “better than” aversive value to context B+0.3 mA. In line with this hypothesis, conditioning mice to contexts A and B both paired with 0.6 mA (**Figure S4K**) led to similar results, with notably a similar tail rattling during S1 and S2 and a similar distance travelled after shock between S1 and S2 (**Figures S4L** and **S4M**). In addition, we observed similar consequences during the choice test with notably a ΔCPP score not different from 0 and similar freezing in A0.6, bridge and B0.6 (**Figures S4N** and **S4O**). We thus propose that aPVT D2+ neurons are critical to compare the relative aversiveness between the two experiences during learning, thereby contributing to the encoding of relative aversive value and triggering appropriate threat-related behavior.

**Figure 4.**
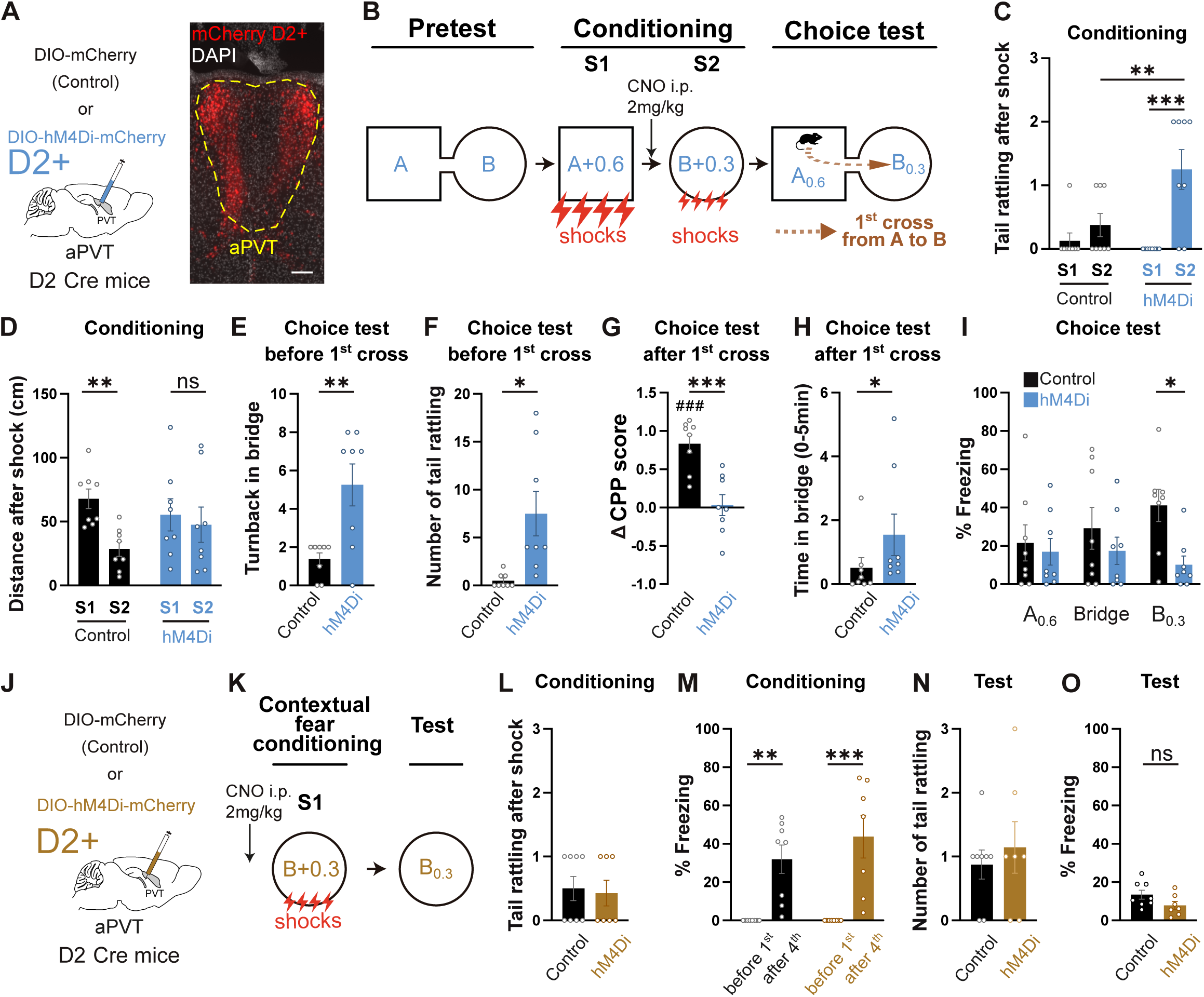

To address whether D2+ neurons are specifically required for relative aversive value coding, but not for coding absolute aversive value, we conducted two sets of experiments in which mice could not generalize threat from a prior aversive experience. In the first experiment, mice were subjected to a contextual fear conditioning task (four shocks of 0.3 mA in context B) (**Figures 4J** and **4K**). We found that silencing aPVT D2+ neurons during contextual fear conditioning of context B+0.3 mA, did not alter the aversiveness of the experience as no changes in freezing nor tail rattling during conditioning and test were observed (**Figures 4L-O**). In a second experiment, mice were subjected to an absolute aversive protocol (no shock in context A followed by four shocks of 0.3 mA in context B; **Figure S4P- Q**). Again, silencing aPVT D2+ neurons during conditioning of context B+0.3 mA, did not change mouse behavior (**Figures S4R-T**). These data are consistent with a specific role for aPVT D2+ neurons to encode relative aversive value during learning.

These results however do not rule out a putative role of aPVT D2- (minus) neurons for the encoding of relative aversive value, considering their role in arousal and responses to stimuli saliency ^33^, and their increased cFos expression during S2 (although significantly lower than aPVT D2+ neurons; **Figure 3D**). To tackle this question, we employed a viral strategy to specifically target and inactivate aPVT D2- neurons (AAV-FLEx-OFF-hM4Di-mCitrine/mCherry with corresponding control viruses based on ^51^; **Figure 5A, Figure S5A** and methods). We first tested the efficiency of the vector by performing *ex vivo* whole-cell recordings in control- and hM4Di-D2- neurons of the aPVT, identified by their fluorescence (**Figure S5A)**. Our data confirmed that bath application of 10□M CNO to aPVT brain sections expressing inhibitory DREADD (hM4Di) hyperpolarizes D2- neurons by drastically decreasing spike frequency. By contrast, bath application of CNO was ineffective on aPVT D2- neurons transduced with the control virus (**Figures S5B**). We also confirmed the specificity of the FLEx-OFF viral vector to aPVT D2- neurons, with only 3% overlap with D2+ neurons (**Figures S5C** and **S5D**). We next injected CNO 30 min before S2 in order to inactivate aPVT D2- neurons during S2 of the aversive task (**Figure 5B**). While hM4Di mice froze slightly less than control before first shock in S2 (**Figures S5E)**, other behavioral parameters previously described remained unchanged during conditioning (including tail rattling, **Figures 5C-D** and **Figures S5F**) nor during choice test (**Figures 5E-I** and **Figures S5G-H)**. Therefore, our results strongly suggest that inactivated aPVT D2+, but not D2- neurons, enhance the relative aversiveness of a threat.

**Figure 5.**
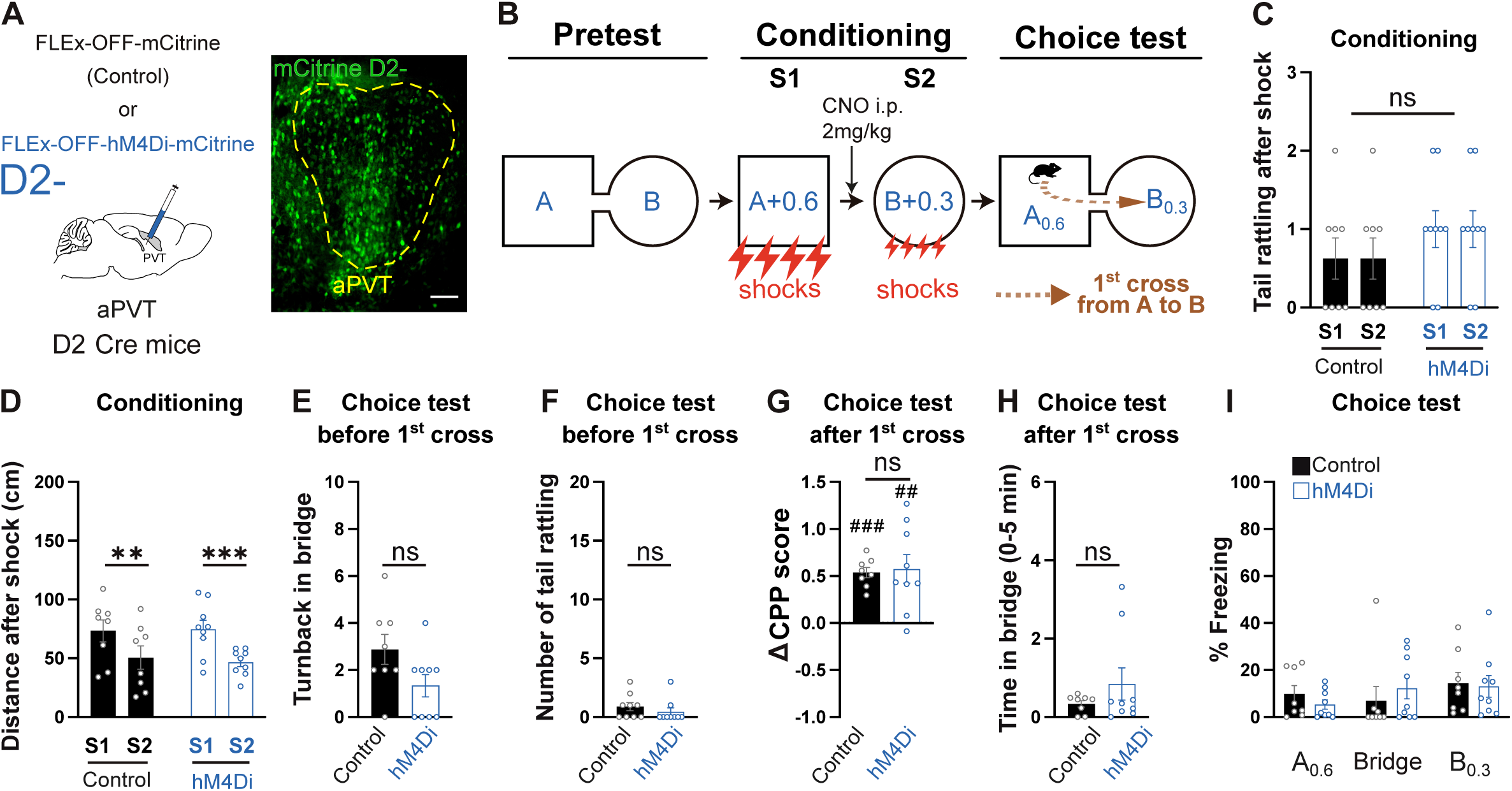

Together, we demonstrated that only aPVT D2+ neurons are essential for the encoding of relative aversive value during learning to enable appropriate relative value-based choice.

### aPVT D2+ neurons dampen relative aversiveness of a threat

To further test our hypothesis that inactivating aPVT D2+ neurons enhances the relative aversive value of the threat-associated context, we conditioned control and hM4Di mice to associate contexts A and B with identical low-intensity shocks (0.3 mA) and administered CNO 30 min before S2 (**Figure 6A**). Confirming our previous findings, silencing aPVT D2+ neurons significantly increased the number of tail rattling post-shock during S2 (**Figure 6B**) without altering the distance travelled after shock (**Figure S6A**). Furthermore, and as previously shown in **Figure S4D**, such pharmacogenetic intervention did not change freezing behavior before the first (**Figure S6B**) or after the fourth shock of each conditioning session S1 and S2 (**Figure 6C**). As a consequence, we found that hM4Di mice first placed in A0.3 during the choice test displayed more turnbacks in the bridge zone before their first cross to context B0.3 (**Figure 6D**), with a similar number of tail rattling (**Figure S6C).** Other behavioral parameters were preserved (first latency, **Figure 6E**, time spent in the bridge zone before, **Figure 6F**, and after the first cross, **Figure S6D** and freezing during choice test, **Figure 6G**). However, as shown in **Figure 4G** during choice test, hM4Di mice did not prefer either of the two contexts (ΔCPP score not different from 0; **Figures 6H** and **S6E**) compared to control mice. This behavior is likely due to the heightened aversiveness of context B0.3 following the inactivation of aPVT D2+ neurons.

**Figure 6.**
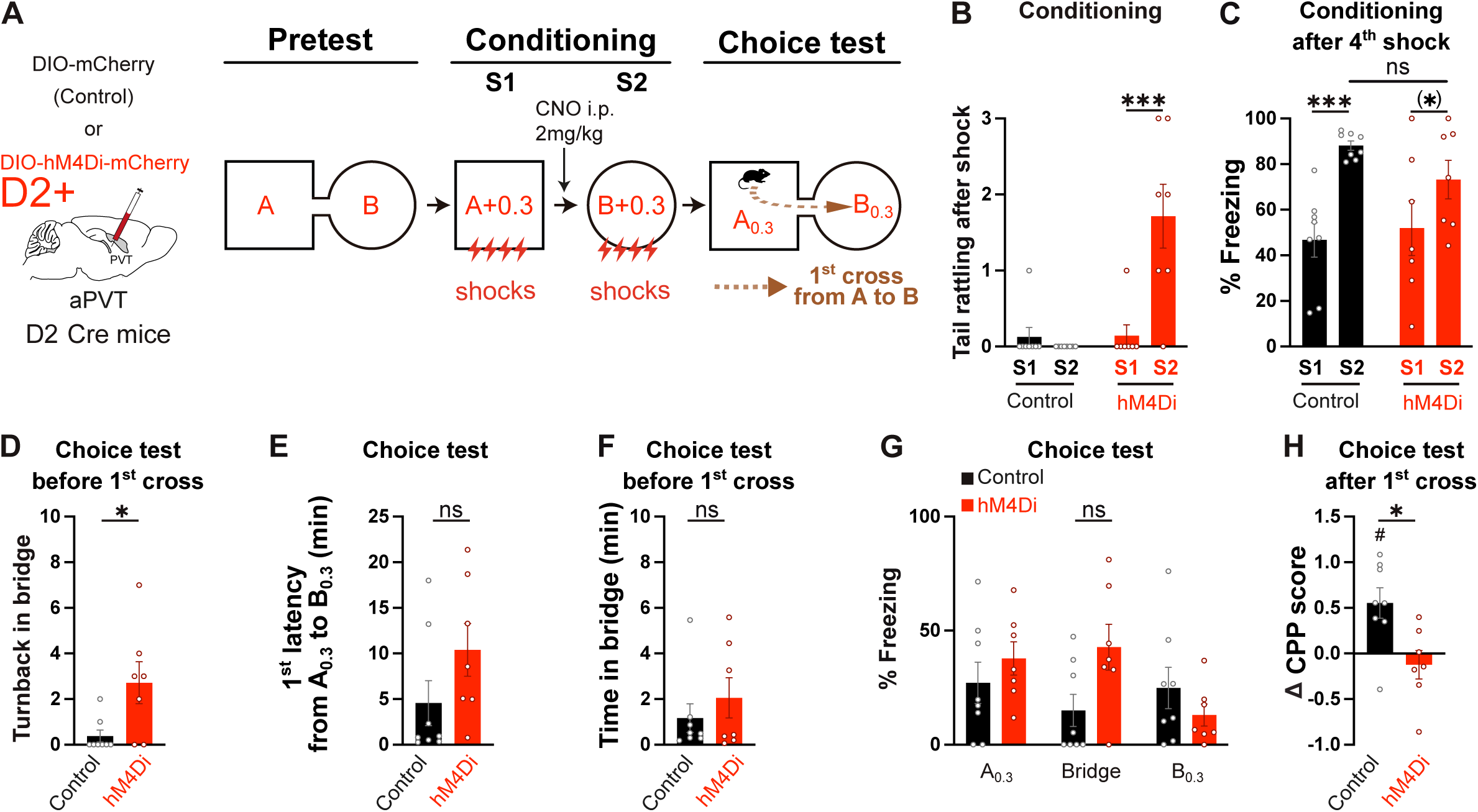

These results combined with those shown in **Figures 4**, **5**, **S4** and **S5** provide strong evidence for a critical role of aPVT D2+ neurons for modulating the relative comparison of aversive experiences during learning and the encoding of relative aversive value, essential information that mice later use to make appropriate value-based choices.

### aPVT D2+, but not D2-, neurons are involved in learned relative safety

PVT neurons modulate the salience of both aversive and appetitive behavior ^28^, a crucial information for both fear and reward learning. ^52,53^ As we found an increased cFos expression in aPVT D2+ neurons in S2 in the safe group (**Figures 3C-3E**), we next asked whether aPVT D2+ neurons also play a role in learned relative safety, as they do for learning relative aversive value. To test our hypothesis, we subjected control and hM4Di mice to the discriminative fear conditioning (**Figure 1A**) and injected CNO 30 min before conditioning S2 in context B+0 mA (**Figure 7A**). In line with our hypothesis, silencing aPVT D2+ neurons did not change threat-related defensive behaviors during conditioning (**Figures 7B- C** and **S7A-B**). During choice test, we also found that threat-predicting behavior such as the number of turnbacks and tail-rattling behavior in the bridge zone before going to the non-shocked B0 context (i.e. before first cross) were also unchanged (**Figures 7D-E**). However, while there were no changes of their first latency to cross from A0.6 to B0 (**Figure 7F**) nor their time spent in the bridge after the first cross (**Figure S7C**), hM4Di mice exhibited a longer latency to go back to the punished context A0.6 (second latency, **Figure 7G**). Importantly, hM4Di mice exhibited a higher percentage of freezing/immobility in the safe context B0 (**Figure 7H**) as they showed a higher preference for the safe context B0 compared to control mice (**Figures S7D** and **7I**). These data indicate that inactivation of aPVT D2+ neurons during S2 increased safety in context B+0 mA, thus strongly increasing the preference towards the safe context B0 during the choice test.

**Figure 7.**
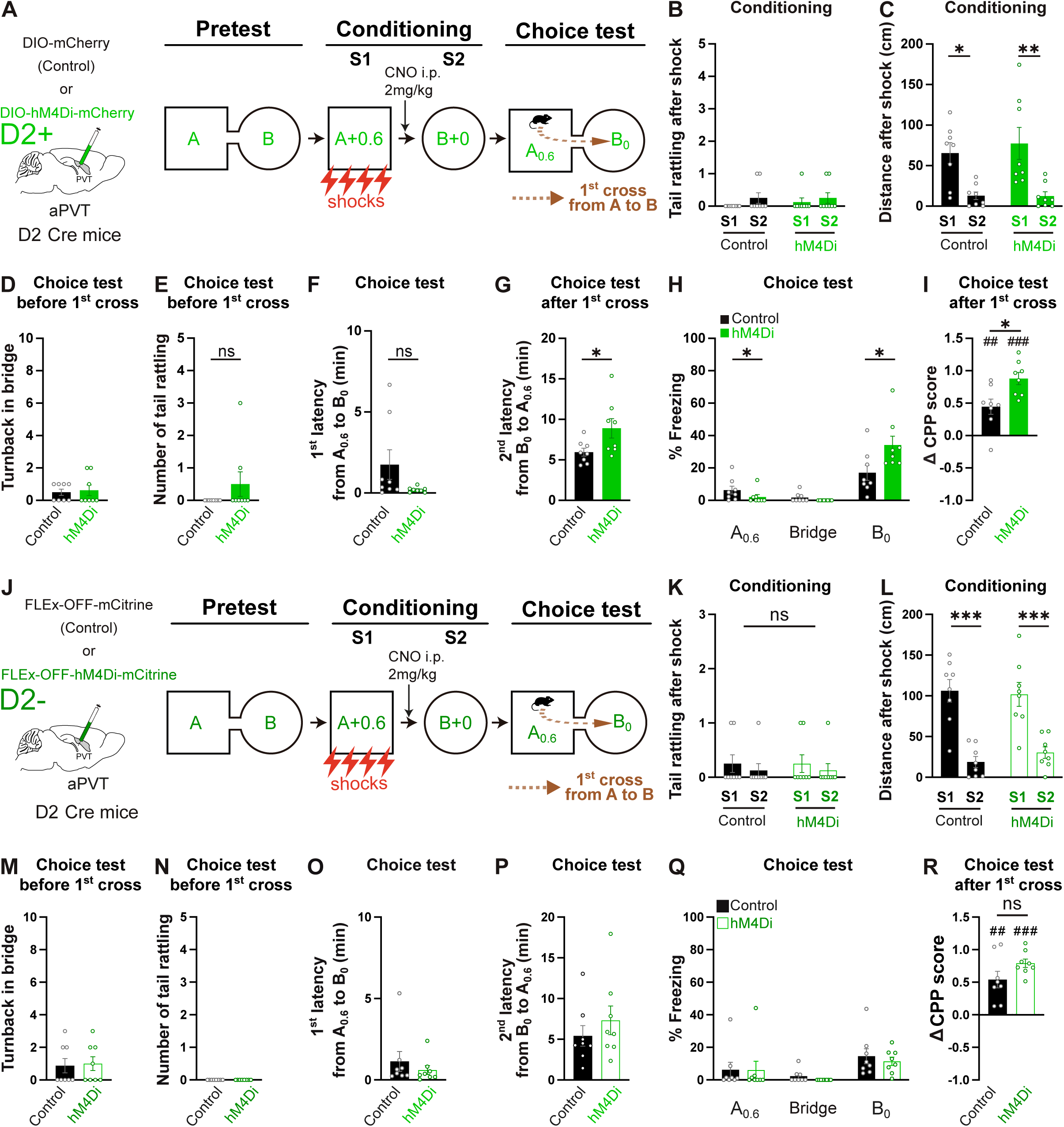

To verify that inactivating aPVT D2+ neurons during the exposure to context B+0 mA did not assign a positive value to this context during conditioning, control and hM4Di mice were subjected to two novel and distinct contexts (C and D contexts, **Figure S7E**) without shock, and CNO was i.p. injected 30 min before S4 in context D+0. As expected, we did not find any change in any mouse behavior during conditioning or choice test (**Figures S7F-O**).

Our results in **Figure 3D** showed an increased cFos expression in aPVT D2- neurons during the encoding of relative safety value, although significantly lower than aPVT D2+ neurons. We therefore asked whether the increased safety we observed reflects the selective inhibition of aPVT D2+ neurons, rather than an overall role of aPVT. To test this hypothesis, aPVT D2- neurons were inactivated during S2 using the same viral strategy as before (**Figures 5** and **7J**). While we found a decreased freezing in S2 for the hM4Di group compared to control mice (**Figure S7P**, as in **Figure S5E**), our results demonstrated that this manipulation had no effect on the capacity of mice to learn a safety value in S2 during conditioning nor choosing to stay in the safe context B0 during choice test (**Figures 7K-R** and **S7 Q-S**).

Together, these findings strongly indicate that aPVT D2+, but nor D2-, neurons modulate learned relative safety during conditioning markedly influencing ensuing value-based choice behavior.

## Discussion

Our study reveals a critical role of aPVT D2+ neurons for learning relative aversive value and learned safety to promote appropriate value-based choices. Using a novel conditioned place preference task, we found that distinct behavioral signatures are associated with learning relative aversive value and learned safety, and related value-based choices. We then demonstrated that learning relative aversive value and learned safety preferentially recruit D2+ neurons in the aPVT compared to that of pPVT. Moreover, inactivating aPVT D2+, but not D2-, neurons at the moment of the comparison between previous aversive and actual aversive/neutral experiences heightened the aversiveness/safeness of the experience. Such enhancement modified learning and then, the selection of subsequent behavior when mice had to choose between the two available options. Importantly, aPVT D2+ neurons modulate relative, but not absolute, aversive value coding, since that in the absence of a prior aversive experience, their pharmacogenetic inactivation did not alter the aversiveness of the ongoing experience. This work provides key findings that improve our understanding of how context-dependent relative (aversive and safe) values are encoded in the brain.

### aPVT D2+, but not D2-, neurons are key modulators of relative aversive value learning

Extensive work focused on brain mechanisms for relative appetitive learning related to reward and gain. ^11,54,54,55,55–65^ Surprisingly, the neural circuits underling relative aversive learning and choice remain poorly understood. Here, we designed a novel 1-day conditioned place preference task in which mice learn the association of two distinct contexts with different shock intensities, the first context paired with 0.6 mA and a second context with less intense 0.3 mA shocks. Later, mice are confronted to a choice between the two contexts. Our data showed that this relative aversive task is associated with increased freezing, emergence of tail rattling and decreased time in the center of the context during the second conditioning session. These key behavioural features were restricted to relative aversive value learning, since they were absent in the absence of prior aversive experience. Strikingly, tail rattling is a behavior typically associated with aggression in male mice before fighting. ^66^ In our set of experiments, tail rattling may therefore be associated with a negative emotional state and a reaction against a possible threat, occurring during learning (i.e. immediately after a negative shock event) but also during memory retrieval (choice test) when mice can make predictions of a possible threat. Previous work has suggested a role of PVT for modulating the value and salience of stimuli. ^28,33^ Moreover, previous studies highlighted a crucial role of the PVT during motivational conflict scenarios where both approach and defensive behaviors compete. ^28,37,48,67^ We found that pharmacogenetic inactivation of aPVT D2+, but not D2-, neurons enhances the aversiveness of the ongoing aversive experience demonstrated by both increased tail-rattling and running immediately after each shock. Since tail rattling behavior may increase visual and auditory saliency of the mouse to predators ^43^, our data suggest that aPVT D2+ neuronal inactivation caused a shift in the defensive behavioral strategy, favoring enhancement of active threat responses.

Our results also revealed that during the relative choice between the two previously punished contexts, mice increased time spent and numerous turnbacks when located in the bridge zone. Such behavior is reminiscent of the “Vicarious Trial and Error” (VTE) phenomenon ^44,68^ observed at decision point when animals pause and move their head back and forth to “scan” and evaluate the possible outcomes before making a choice between two or more options with uncertain outcomes. ^45^ Importantly, VTE behavior is preferentially exhibited in hippocampal-dependent tasks and during complex decision making processes. ^69–71^ Given the critical role of contextual information in our new relative task and the observed PVT-hippocampus anatomo-functional connectivity ^23^, our newly designed relative aversive task could serve to expand our knowledge of the neuronal circuits underlying VTE behavior and value-based decision-making processes.

### aPVT D2+ neurons modulate predictive signals for learning aversive value and learned safety

The PVT is widely recognized as signaling aversive states and modulating fear-related behaviors ^23,28,33,36,38,41,72–74^, as well as appetitive associative learning ^28,75^, and reward-seeking behavior. ^76–78^

Moreover, PVT neurons are activated by cues or contexts previously associated with positive or negative emotional outcomes. ^23,36–38,73,76,79–84^ In the safe condition, in which the first context is paired with 0.6 mA shocks followed by a second context in which no shocks are delivered (a differential/discriminative conditioning), animals likely acquire a learned safety. ^4,12–14^ Our cFos staining revealed that the aPVT is more recruited than pPVT by both a second context paired with a weaker shock (relative aversive) or the absence of shock (learned safety). As mice likely generalize expected negative prediction from a previous aversive experience to learn relative aversive value or safety, our results are reminiscent of previous reports showing that the omission of expected reward activates the aPVT, but not the pPVT. ^76^ The aPVT may therefore be strongly involved in generating predictive value signals crucial for the encoding of relative aversive/safety values requiring the reward system. To compute these predictive value signals, the PVT receives inputs from regions such as the infralimbic, prelimbic cortex ^27,85–88^ and ventral hippocampus ^89,90^, which convey information about motivational state and arousal, respectively. To modulate predictive value signals, the PVT projects to the ventral tegmental area (VTA) and substantia nigra pars compacta ^91^ where computation of prediction error (comparison between predicted and actual information) is crucial to signal correct value to sensory experience during learning.^92,93^ Notably, VTA dopaminergic neurons can regulate the salience of threat signals. ^94^ As the reward system is involved in both relative aversive value ^6,7^ and learned safety ^14,16,17,95^, the aPVT could modulate the computation of a relative positive value signals in the VTA. The aPVT also projects to regions modulating approach-avoidance behaviors such as prefrontal cortex ^33,96^ and ventral subiculum ^96^, extended amygdala ^96,97^, nucleus accumbens (NAc) shell ^26,96,98^ and NAc core. Since the NAc shell region integrates dopaminergic inputs from the VTA and (indirect) glutamatergic inputs from the PVT ^99,100^, inactivating the PVT-NAc pathway originating from aPVT D2+ neurons ^101^ could attenuate dopaminergic release in the NAc. Dopamine in the NAc is increased in response to safety cues^14^, and recent work reveals that the PVT is required for this dopaminergic safety signal. ^102^

Inactivating the aPVT D2+ neurons could lead to an aberrant assignment of value to neutral stimuli ^103^ and therefore disrupt the ability to learn relative aversive and safety values. Unlike the pPVT, the aPVT also receives major hippocampal subiculum projections. ^27^ During the relative comparison of (aversive/safe) experiences, contextual information from the subiculum could be instrumental for the generalization of threat prediction that aPVT D2+ neurons use to modulate reward VTA and/or NAc circuits to enable the encoding of relative aversive/safe value.

Unlike innate safety towards a novel “naïve” CS, learned safety involves the discrimination between CSs to avoid overgeneralization of a threat. Although several behavioral paradigms have been developed to examine learned safety ^14^, most of the literature has focused on tasks where the animals tested cannot escape a potentially dangerous situation, thereby restricting their repertoire of fear related behaviors. This has led to safety been classically measured through freezing behavior ^42,104,105^. However, paradigms beyond instrumental tasks ^106,107^ can incorporate safety approaching measurements, and future studies should include measurements of safety such as the active approach of a safe compartment as seen in conditioned place preference/avoidance paradigms such as ours.

Furthermore, our results revealed a preferential activation of aPVT D2+ neurons during learning relative aversive value or safety, while paradoxically we and others found that D2+ neurons are preferentially located in the pPVT. ^33,49^ Importantly, generalized threat prediction, involved here for learning relative aversive value or learned safety, also involve the activation of D2 type receptors in the extended amygdala substructures i.e., the central amygdala and the bed nucleus of the stria terminalis that have been shown to contribute to anxiety-like behaviors. ^108^ Of note, aPVT D2+ neuronal activation was tightly linked with the time spent in the bridge during choice test, a crucial behavioral hallmark of the decision-making process between different options. Indeed, the less recruited aPVT D2+ neurons, but not pPVT, the more animals spent time in the bridge in the relative aversive, but not safe, condition. Consistent with this observation, we found that the selective inactivation of aPVT D2+ neurons increased the time spent in the bridge in the relatively aversive, but not the safe, condition thus preventing mice from selecting the most appropriate behavior when facing a complex decision between two aversive situations.

### aPVT D2+ neuronal modulation of salience vs behavioral disinhibition

Emotionally salient events, regardless whether positive or negative, are encoded and remembered with greater accuracy compared to emotionally neutral ones. ^109–113^ The PVT is involved in arousal ^33,114,115^ and can encode salience of both conditioned and unconditioned stimuli. ^28^ Prior work revealed the existence of two subpopulations of PVT neurons that form molecularly, anatomically and functionally distinct subpopulations. ^33,49^ For instance, D2+ neurons (type I neurons) send projections to the prelimbic cortex and are sensitive to aversive stimuli. ^33,39^ By contrast, D2- neurons (type II neurons) send projections to the infralimbic cortex and are modulated by stimulus salience. ^33^ However, given the preferential anatomical distribution of D2+ neurons in the pPVT ^33,35,49,101^, most studies neglected the contribution of D2+ neurons located in the aPVT. Further studies will therefore be required to disentangle the role of the projections originating from D2+ or D2- neurons of the aPVT (targeting the amygdala or NAc ^33^). In this sense, our work not only sheds light on the role of aPVT D2+ neurons but also, highlights, more broadly, the profound influence that even a relatively small neuronal population can exert on relative value-based learning.

Our results showed that the inactivation of aPVT D2+, but not D2-, neurons enhances the relative aversiveness or safeness of the ongoing experience, possibly by increasing its salience. Yet, it remains unclear whether the conditioned stimulus (here, the context) or the outcome (shock or no shock delivered) become more salient. In line with the latter option, such manipulation for the relative aversive task increased tail rattling behavior immediately after the shock delivery during conditioning, but not general freezing. Consistently, a recent study identified that tail rattling, but not freezing, behavior positively scales with arousal level. ^43^ However, the precise neural mechanisms through which aPVT neurons contribute to the expression of active defensive behavior such as tail rattling remain to be defined. Inactivation of aPVT D2+ neurons also increased the number of turnbacks and time in the bridge zone (i.e., hesitation) during the choice test, thus leading to the absence of preference for the less punished context. Importantly, the same inactivation procedure during the safe conditioning did not affect tail rattling behavior nor the time in the bridge zone, but it led to an increased preference for the safe context. The efficiency of the value-based choices could therefore be influenced by the salience of the experience and the computation and comparison of values assigned at the time of learning. ^18^ Inactivation of aPVT D2+ neurons could have led to an incorrect value assignment during the context-outcome association, thus favoring the contribution of D2- neurons for the attribution of salience to the ongoing experience. ^33,82^ This could hamper (for the relative aversive task) or favor (for learned safety) the ability of animals to distinguish between the options associated with each context. Inactivating aPVT D2+ neurons could therefore enhance the formation of memory of any otherwise low-salient stimuli. An alternative explanation could be that aPVT D2+ neurons provide a specific function in behavioral disinhibition. ^48,76,116–119^ Then, inactivation of aPVT D2+ neurons during conditioning S2 could unleash context-dependent behavior such as active defensive (tail rattling) behavior during relative comparison of aversive experiences or decreased freezing and increased preference for the safe context. Indeed, aPVT D2+ neurons could exert context-dependent behavioral disinhibition by modulating the salience of contextual cues/information, which allows mice to detect the aversiveness or safeness of an experience to disinhibit specific and adapted behaviors. However, testing these hypotheses will require further investigation.

Unlike the opposite effect observed with D2+ neuronal inhibition over relative safety and aversive value learning, D2- neurons seem to be engaged (increased cFos expression) in a similar manner independently of whether the animal is comparing between two aversive experiences or an aversive and a neutral one. Beas et al 2024 found a decreased activity of aPVT D2- neurons as animals approach a reward.^120^ Thus, inhibiting aPVT D2- neurons in our tasks may have transiently reduced generalized fear expression (i.e., decreased freezing to contextual cues) without affecting the mouse’s ability to learn relative aversive/safety value.

Overall, our findings extend current knowledge about the role of PVT as an integrator of emotional, contextual and sensory information to guide appropriate behavior. This study provides new evidence regarding the role of aPVT D2+ neurons for contextual relative aversive/safety learning that supports appropriate value-based decisions. A better understanding of the neuronal circuits and mechanisms responsible for the balance between value information, salience and behavioral disinhibition is therefore critical for deciphering motivated behaviors. Indeed, an excessive stimulus evaluation and/or salience processing during traumatic events (that could involve the PVT) might lead to associative learning impairment and contribute to the genesis of anxiety disorders such as PTSD. ^121,122^

## Resource availability

Further information and requests for resources and reagents should be directed to, and will be fulfilled by, the corresponding authors, Emmanuel Perisse and Stéphanie Trouche.

## Materials availability

All original reagents presented in this study are available from the Lead Contacts upon request.

## Data and code availability

All code used in this paper using Fiji will be shared by the Lead Contacts upon request.

## Methods

### Animals

We used C57BL/6J (Charles River Laboratories, France) and three transgenic heterozygous adult male mouse lines: D2 Cre mice (STOCK Tg(*Drd2* cre) ER44Gsat/Mmucd, provided by E. Valjent), D2 Cre:Ribotag mice ^123^ (provided by E. Valjent) and D2 Cre:Ai14 mice, that were generated by crossing Drd2 Cre mice with Ai14 mice (B6.Cg-*Gt(ROSA)26Sor^tm^*^14^*^(CAG-tdTomato)Hze^*/J, stock #007914). The latter transgenic mouse line expressed the red fluorescent protein double-tomato (tdTom) under endogenous regulatory elements of the D2 receptor. All transgenic mice were maintained on a C57BL/6J background. Animals (3 to 5-month-old mice) were housed with their littermates and were singly housed in their home cage after the surgical procedure on a 12-h light/dark cycle (7 a.m. to 7 p.m.) in stable temperature (22°C) and humidity (60%) conditions with free access to food and water in a dedicated housing room. Mice were randomly assigned to behavioral groups. All experimental procedures were conducted in agreement with the guidelines of the French Agriculture and Forestry Ministry for handling animals (authorization number/license A3417241) and approved by the local and national ethic committees (authorizations APAFIS #23355).

### Immunofluorescent staining and c-Fos analysis

Mice (D2 Cre:Ai14 or D2 Cre:Ribotag, see **Figures 3** and **S3**) were anaesthetized and transcardially perfused with PBS followed by 4% PFA in PBS solution 90 min after the second session of conditioning of the CPP task. Brains were extracted and kept in 4% PFA for at least 24 h before slicing using the vibratome and coronal sections (30μm thick) were stored in PBS-azide at 4°C. For the immunohistochemistry procedures, free-floating sections were rinsed extensively in PBS with 0.25% Triton X-100 (PBS-T) and blocked for 1 h at room temperature in PBS-T with 10% normal donkey serum (NDS). An average of 3.5 sections per animal of either aPVT or pPVT were used. Sections were then incubated overnight at room temperature with primary antibodies diluted in 3% NDS blocking solution. The primary antibodies used were: anti-RFP (MBL, rabbit, 1/2000, #PM005), *c-Fos* antibody (Synaptic Systems, rat, 1/5000, #226017), anti-HA (rabbit, 1/500, Rockland #600-401-384) and anti-GFP (chicken, 1/2000, #GFP-1010). After three 15 min washes using PBS, sections were incubated for 2h30 at room temperature in PBS containing a mixture of secondary antibodies. Secondary antibodies used were: donkey 488-coupled anti-rabbit (Jackson Laboratory, 1/500, #711-545-152), donkey Cy3-coupled anti-rabbit (Jackson Laboratory, 1/1000, #711-165-152), donkey Cy5-coupled anti-rat (Jackson Laboratory, 1/500, #712-605-150), donkey 488-coupled anti-rat (Jackson Laboratory, 1/500, #712-545- 153) and donkey Cy3-coupled anti-rat (Jackson Laboratory, 1/1000, #712-165-153). After three 15 min washes in PBS, sections were mounted on slides and stored at 4°C. Images were taken using a ZEISS Apotome microscope at 16bits and either 10X or 20X. cFos expression was examined in D2 tagged cells using the D2 Cre:ai14 mice or D2 Cre:Ribotag (in a balanced manner by group) using ImageJ based automated analysis. For each section, a region of interest was assigned for both aPVT and pPVT and a threshold was selected to count both the amount of c-Fos+ cells, D2+ cells and c-Fos+ D2+ colocalization. Sections from aPVT were taken at AP:-0.22 to -1.22 while the pPVT were taken at AP:-1.34 to -2.18.

### *In vivo* pharmacogenetic inhibition

For *in vivo* pharmacogenetic (Designer Receptors Exclusively Activated by Designer Drugs, DREADD), inhibition in CPP and contextual fear conditioning (CFC) experiments, Clozapine N-Oxide (2 mg/kg CNO, Enzo Life Science, #BML-NS105-0025) was dissolved in 0.9% NaCl saline solution and intraperitoneally (i.p.) injected 30 min before the conditioning session (for CFC) or the second conditioning session (for CPP).

### Stereotaxic viral injections

The following adeno-associated viral vectors (AAV) were used: AAV2/1-hSyn-DIO-hM4D(Gi)-mCherry, (hM4Di, 7.1E12 vg/mL, #44362 Addgene), AAV2/1-hsyn-DIO-mCherry (control, 1.7E13 vg/mL, #50459 Addgene), AAV2/9-hSyn-FLEx-OFF-hM4D(Gi)-mCherry (hM4Di, 1.1E14 vg/mL, Plasmid #161579 Addgene), AAV2/9-hSyn-FLEx-OFF-mCherry (control, 6.1E14 vg/mL, designed in house from Plasmid #161579 Addgene), AAV2/9-hSyn-FLEx-OFF-hM4D(Gi)-mCitrine (hM4Di, 1.8E15 vg/mL, designed in house from Plasmid #161579 Addgene), AAV2/9-hSyn-FLEx-OFF-mCitrine (control, 1.9E15 vg/mL, designed in house from Plasmid #161579 Addgene). Mice were anaesthetized with isoflurane (0.5– 3.5% oxygen gas mixture 1 l/min) and buprenorphine (0.1mg per kg of body weight) for stereotaxic injections. Viral injections were targeted unilaterally to aPVT (coordinates: AP -0.40 mm; ML -0.60 mm; DV -3.25 mm, 10° angle; 200 nl per site). The viral vectors were delivered at a rate of 100nl/min using a glass micropipette lowered to the target site and that remained for 5 min after injection before it was withdrawn. Mice were tested for behavior 4-5 weeks after viral injections. At least 7 days before testing, animals were singled caged to prepare them for the behavioral experiment and trained with mock i.p. injection.

### Behavioral procedures

#### Conditioned Place Preference experiments

We developed a 1-day aversive task based on a previously published conditioned place preference (CPP) task. ^46,47^ The apparatus contained two electric grids on the floor of each context and consists of two square-walled (46 cm × 46 cm × 30 cm) contexts with distinct inside building block configurations. The two contexts were connected via a bridge (8-cm length, 7-cm width, 30cm height) during both pretest and test sessions. The access to the bridge was blocked during each conditioning session. All mice were handled for at least 3 days before the first experiment and, when experiments involved viral injections, habituated to the injection procedure by mock injections during these days.

During pretest, mice freely explored the two distinct contexts (e.g. one square and one circle) for 15 min and their baseline (spontaneous) preference for one of the two contexts was determined. Next, mice were subjected to a first 10 min-conditioning session (S1), in either no shock (0 mA, "A+0") or 4 shocks (0.6 mA or 0.3 mA of intensity, at 198 s, 278 s, 358 s and 438 s, 2-s duration each, as in ^124^) were delivered in their most preferred context (Context A, "A+0.6" or "A+0.3"). Mice returned to their home cage at the end of session S1 for 30 min. During the second 10 min-conditioning session S2, mice received, in their least preferred enclosure (Context B), either no shock (0 mA, "B+0") or 4 shocks (of 0.6 mA for "B+0.6" or 0.3 mA for "B+0.3" at 198 s, 278 s, 358 s and 438 s, of 2-s duration each). The control group (0 vs 0) went through the two sessions S1 and S2 without receiving any shock. During the choice-test session 1 hour later without any shock delivered, mice were placed in their most preferred context A (as defined during pretest) and were free to visit the two enclosures for 15 min after their first cross from context A to context B. For **Figure S4 P-T**, the mouse was placed in the non-preferred (punished) context D during the choice test, and was free to visit the two enclosures for 15 min after their first cross from context D to context C. Mice that displayed more than 75% preference during pretest were excluded for the rest of the procedure.

#### Contextual fear conditioni3ng experiments

A separated group of mice shown in **Figure 4** was subjected to classical contextual fear conditioning. During the 10 min-conditioning session (S1), mice received 4 shocks of 0.3 mA each in Context B ("B+0.3" at 198 s, 278 s, 358 s and 438 s, of 2-s duration each). All mice were tested in the same context 1h later without any shock delivered for 15min (retrieval test session).

#### Behavioral analysis

Mice were tracked and several behavioral parameters were measured during pretest, conditioning and choice test. A conditioned place preference score (CPP score) was calculated for each mouse during both pretest and choice test as in ^47^. Mice that did not cross to the non-preferred context B in less than 25 min were excluded from further analysis (i.e. they did not perform the following 15 min choice test).

CPP was calculated as 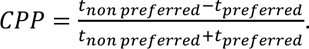 ΔCPP was calculated as the difference between CPP test and CPP pretest. The first latency to cross was the time the animal takes to first cross from the originally preferred context A (where they were first placed) to the originally non-preferred context B. The second latency to cross was the time the animal takes to cross back to the preferred context A. Time in the bridge was measured during the first latency (time in the bridge before first cross) and during the first 5 min after the first cross. Freezing behavior was systematically measured using a custom-made software (Imetronic) and was calculated as the percentage of time for which mice remained immobile for 2 s or more. The percentage of time spent freezing during the entire duration of the pretest and test session were reported. For each conditioning session, the percentage of freezing before the 1^st^ shock (min 1-3) and after the 4^th^ shock (min 9-10) were calculated. For **Figure 1B**, the difference between the total percentage of freezing in S2 minus the total percentage of freezing in S1 was calculated for each group (total percentage of freezing for S1 and S2 used for this calculation are presented in **Figure S1**). All relative measurements (ΔCPP, ΔTime in bridge, etc) were calculated by subtracting choice test behavioral measurements with the corresponding measurement during the pretest. For example, ΔTime in the bridge was calculated as the time spent in the bridge during the first 5 min after crossing during the choice test minus the time spent in the bridge during the first 5 min of the pretest session, ie. Δ0123 451673 = 0123 451673 "ℎ91:3_;.<=>?_ − 0123 451673 #53A3BA_;.<=>?_. We quantified whether the animal engaged in turn back behavior (i.e. when the whole body turns at 180 degree) before 1st crossing from A to B when the animal was in the bridge. Tail rattling was measured as a rapid vibration of the tail either after each shock delivered during conditioning sessions S1 and S2 or before the first cross during the choice test. For experiments shown in **Figure S4**, animals from the 0-0.3 condition were animals that went through CFC one week before. A different set of contexts (C and D) with electric grids. For experiments shown in **Figure S7**, animals from the 0-0 control group were animals that went through the 0.3-0.3 condition one week before. A different set of contexts (C and D) were used in which the floor was plain and without grids. For measuring the distance travelled post shock delivery as done in ^125^, we averaged for each conditioning session the distance travelled for 8 sec at the onset of each shock event (i.e. 198-206 s, 278-286 s, 358-366 s, 438-446 s).

In **Figure 5**, 1 mCherry animal was excluded for not crossing during choice test, 2 hM4Di and 1 mCherry animals were excluded for transduced cells found outside PVT. In **Figure 6**, 1 hM4Di and 1 mCherry animal were excluded for not crossing during choice test. In **Figure 7**, 1 hM4Di animal was excluded for transduced cells found outside PVT.

### Brain slice electrophysiology

Patch-clamp recordings, in whole-cell configuration, were performed on PVT D2-labeled cells in current clamp mode. Briefly, injected D2 Cre mice (n=4 for Cre ON condition and n=4 for Cre OFF condition; for each condition, 2 mCherry and 2 hM4Di mice) were anesthetized and perfused intracardially with ice-cold dissection buffer (25 mM NaHCO3, 1.25 mM, NaH2PO4, 2.5 mM KCl, 0.5 mM CaCL2, 7 mM MgCl2, 25 mM D-glucose, 110 mM choline chloride, 11.6 mM ascorbic acid, 3.1 mM Pyruvic acid maintained in 5% CO2/ 95% O2). Coronal brain slices (350 µm) were cut in dissection buffer using a vibratome (Campden Instruments). Sections were incubated in artificial cerebrospinal fluid (ACSF) containing the following (in mM) 125 NaCl, 2.7 KCl, 11.6 NaHCO3, 0.4 NaH2PO4, 1 MgCl2, 2 CaCl2, 5.6 D-glucose (pH 7.4, osmolarity 305 mOsm) at 32°C for 5 min and then kept in room temperature for at least 1h before recording. For the experiment, slices were superfused continuously at 1.5 ml/min with an ACSF solution. Recordings were obtained using a Multiclamp 700B amplifier (Molecular Devices). Recording electrodes made of borosilicate glass had a resistance of 4-6 MΩ (Warner Instruments, USA) and were filled with K-gluconate solution containing (in mM) 125 K-gluconate, 5 KCl, 10 Hepes, 1 EGTA, 5 Na-phosphocreatine, 0.07 CaCl2, 2 Mg-ATP, 0.4 Na-GTP (pH 7.2, osmolarity 300 mOsm). Only cells with access resistance < 20 MΩ and a change of resistance < 25% over the course of the experiment were analyzed. Data were filtered with a Hum Bug (Quest Scientific, Canada), digitized at 2 kHz (Digidata 1444A, Molecular Devices, Sunnyvale, CA, USA), and acquired using Clampex 10 software (Molecular Devices). Whole cell recordings were made from mCherry aPVT D2+ or D2- neurons visualized with an Olympus microscope (equipped with an mCherry filter). CNO was dissolved in ACSF (10µM, 300 s) and applied in the bath after 5 min baseline recording. The number of action potentials were analyzed before and after CNO application with Clampfit 10 software (Axon Instruments).

### Statistical Analysis

Data were shown as the mean ± SEM (standard error of the mean). GraphPad Prism v10 software was used for statistical analyses. D’Agostino-Pearson omnibus test is used as a test for normality. When this assumption was not met, non-parametric tests were used (see **Table S1**). Student’s t-test (paired or unpaired, 2-sided) was used to compare two sets of data. Multiple comparisons were performed by 1-way or 2-way ANOVA followed by either Sidak’s, Tukey’s, Uncorrected Fisher’s LSD or Dunn’s multiple comparisons were used for *post-hoc* analysis. For comparisons against 0, either one sample t test or Wilcoxon signed rank test (for non-normal data) were used. All statistical comparisons are presented in **Table S1**.

## Acknowledgements

We warmly thank S. Parkes and G. Gangarossa for their valuable comments on the manuscript. We thank M. Verdy for his help with setting up the behavioral task. We thank the vector core facility of Montpellier /Plateforme de Vectorologie de Montpellier, BioCampus, UM-CNRS-INSERM, Montpellier, France, especially C.Lemmers for producing the Cre Off viruses. We thank E. Valjent for sharing the D2 Cre mouse line. This work was supported by CNRS, INSERM, the IGF, the ATIP-Avenir program (EP), the Bettencourt Schueller Foundation (EP), the French Research Agency (ANR-19-CE37-0006, ST and LV; ANR-21-CE16-0015, EP, ST, MD and EK), the CBS2 doctoral school of Montpellier (EK), the Human Frontier Science Program Postdoctoral Fellowship (LT0027-2022-L, MM) and Fyssen Foundation (MM). We acknowledge the imaging facility MRI, member of the France-BioImaging national infrastructure supported by the French National Research Agency (ANR-10-INBS-04, “Investments for the future”). We also thank all the Iexplore staff from IGF, especially M. Dedin, for their help in the maintenance and breeding of mice.

## AUTHOR CONTRIBUTIONS

M.M., E.P. and S.T. conceived the project and designed all the experiments. M.M. and E.K. performed surgical and anatomical experiments. M.M., E.K., L.V., M.D and C.V. performed behavioral experiments. P.T., M.P, E.K. and J.S. performed and analyzed electrophysiological experiments. M.L. and E.K. helped with cloning constructs and viral strategies. The manuscript was written by M.M., E.P. and S.T and commented by all authors. All authors were asked for input on the text.

---

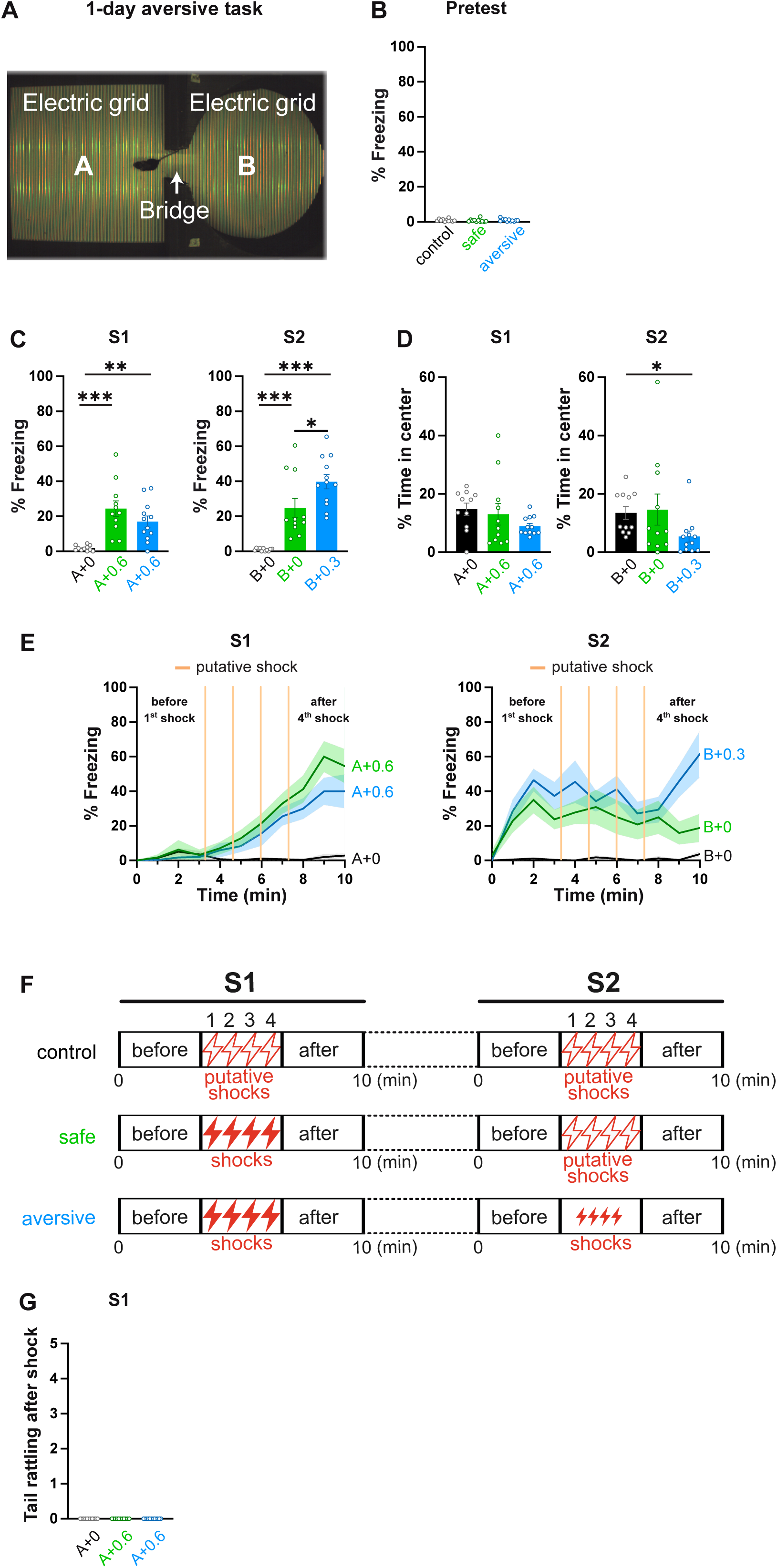

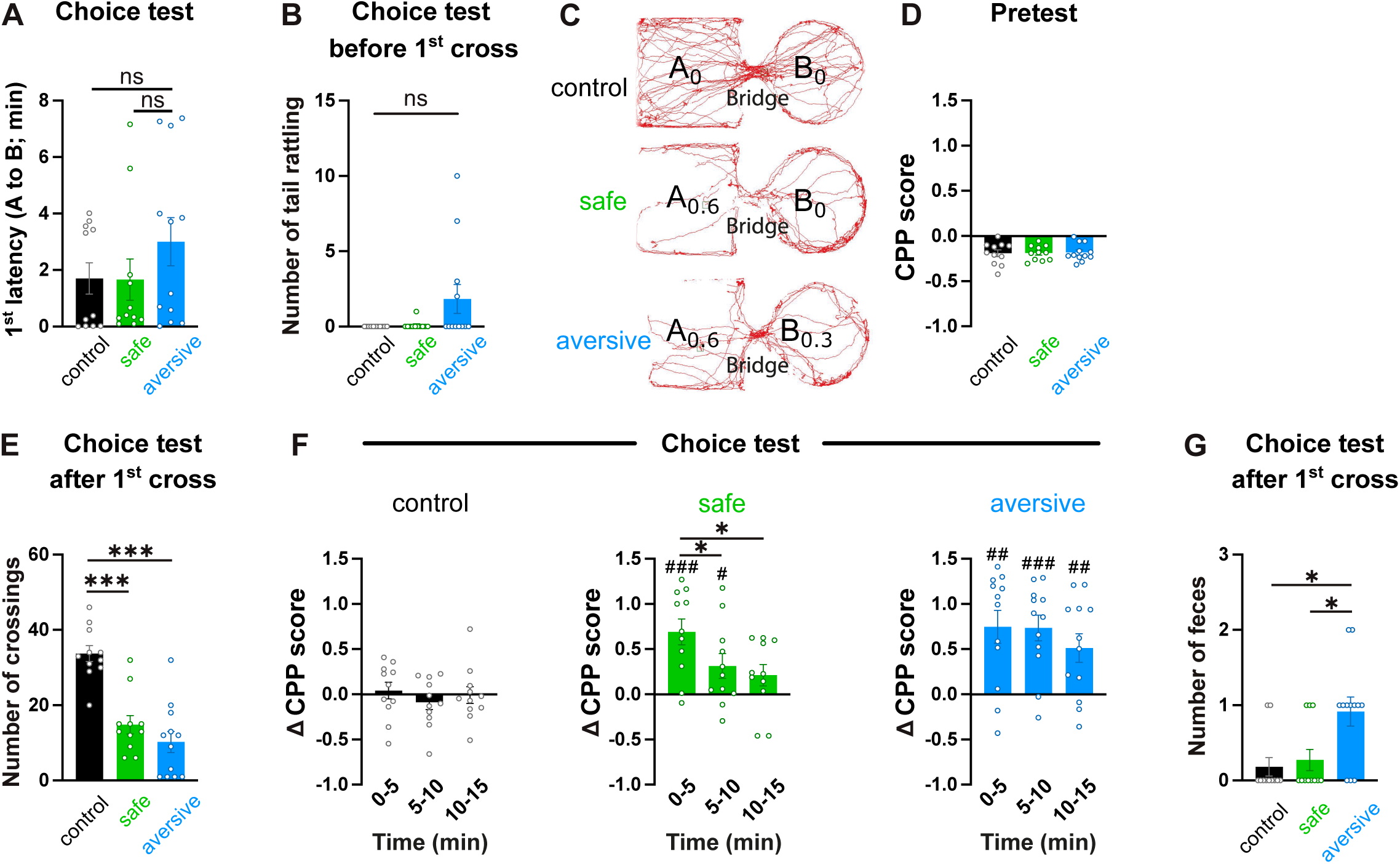

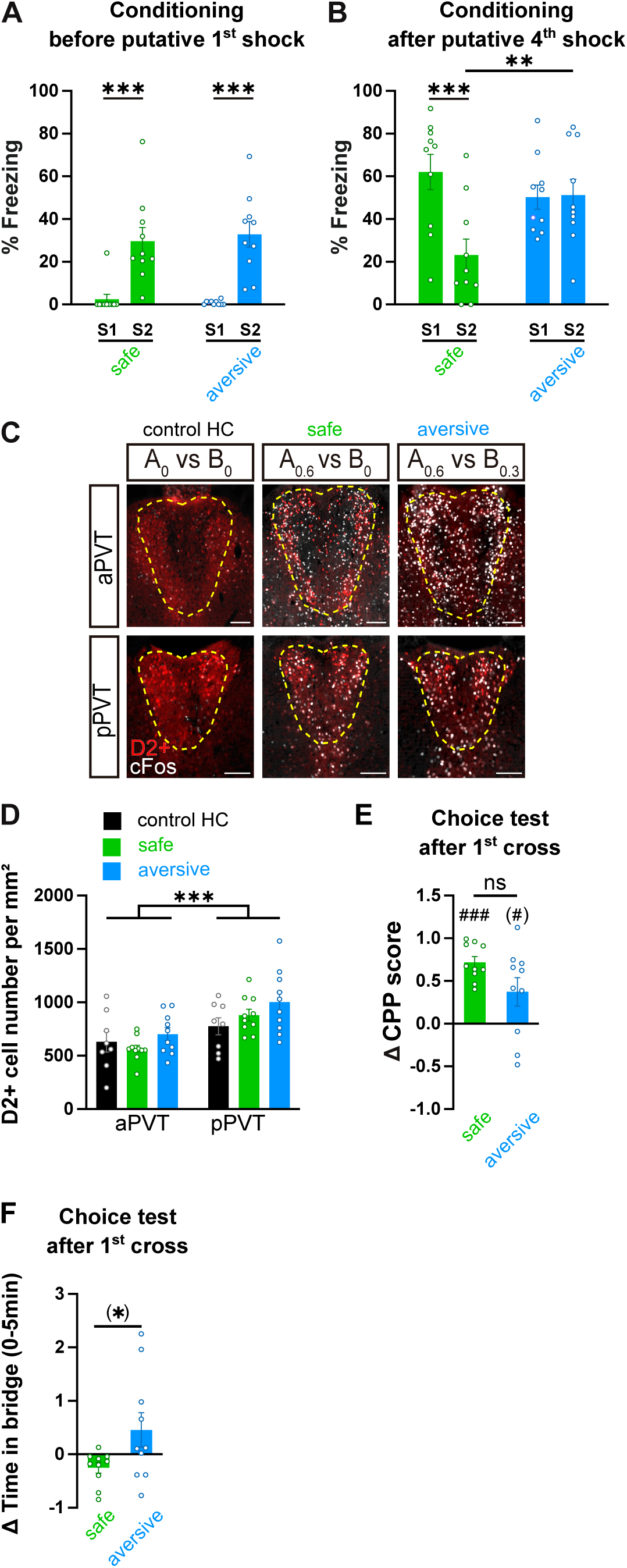

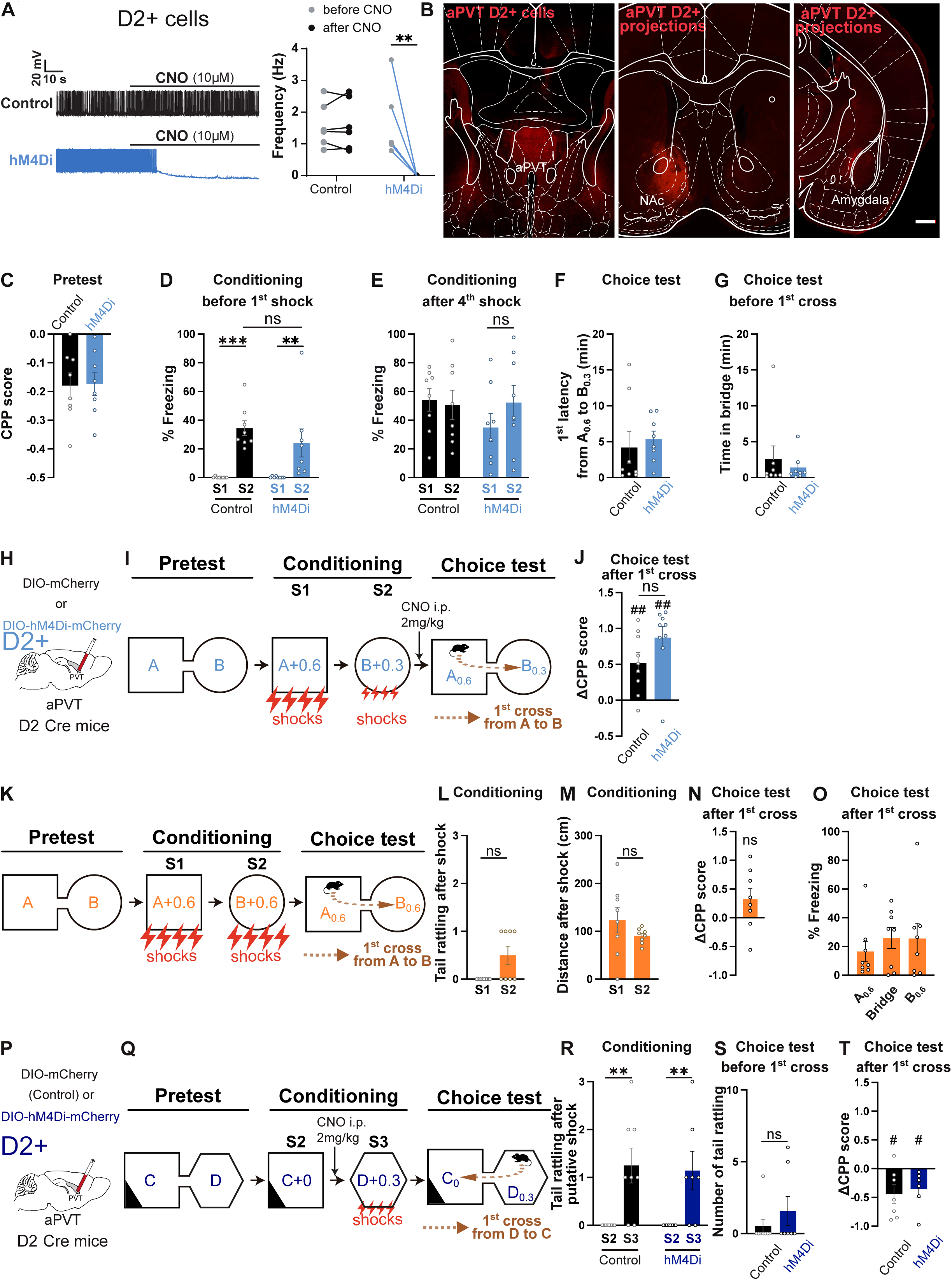

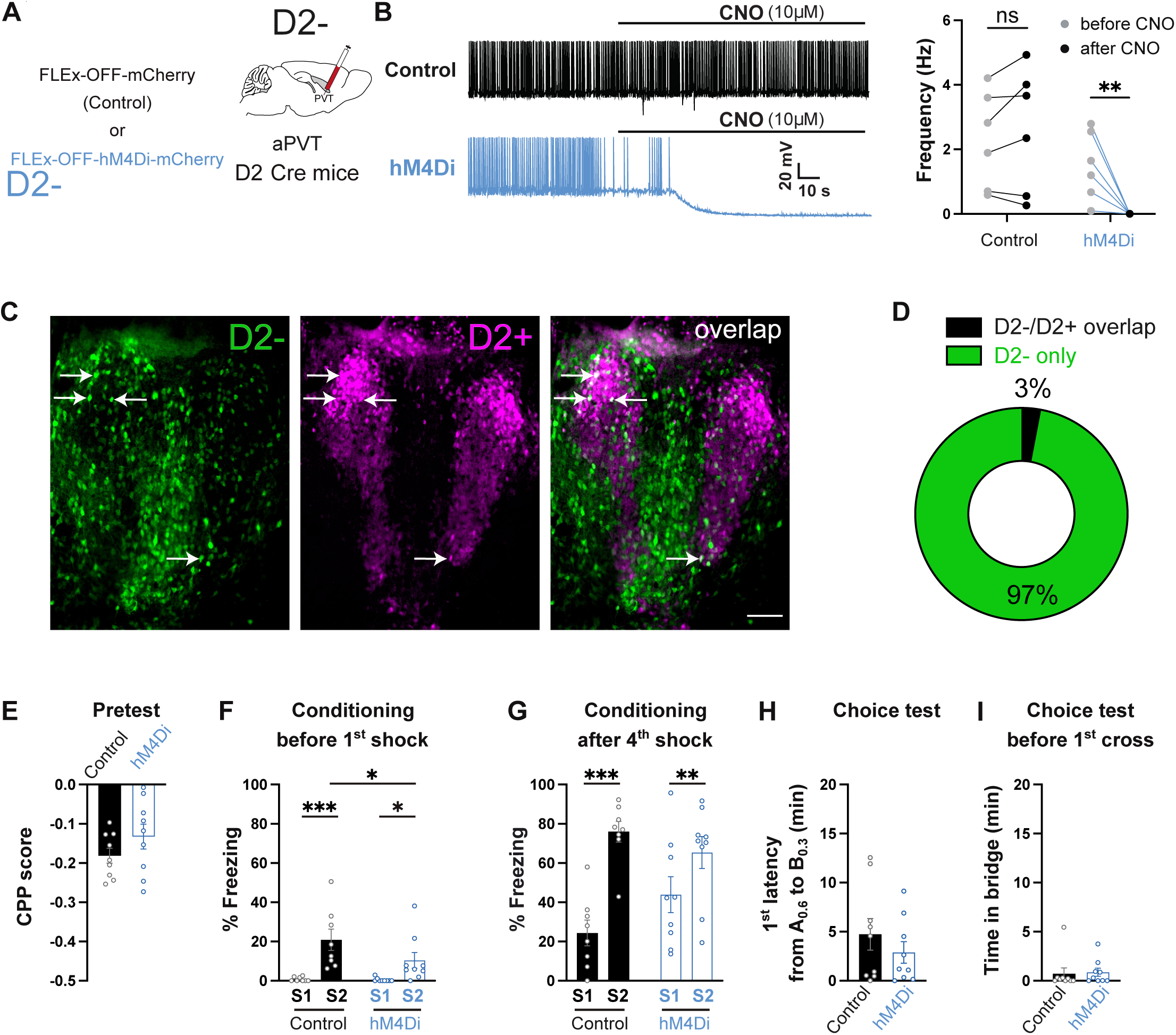

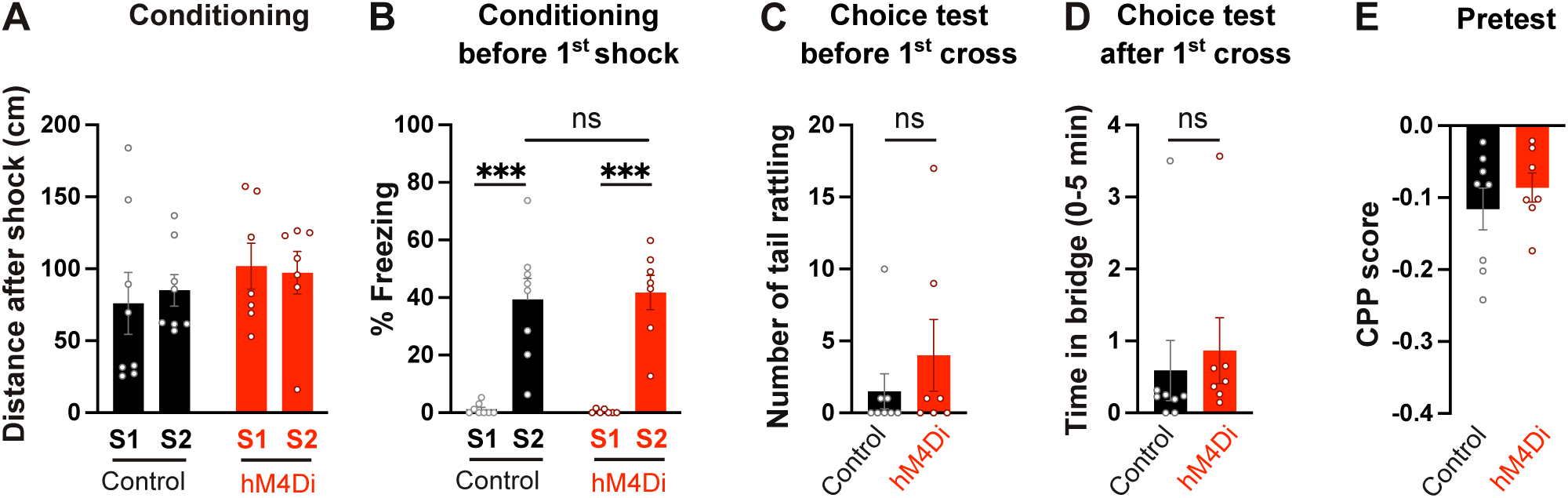

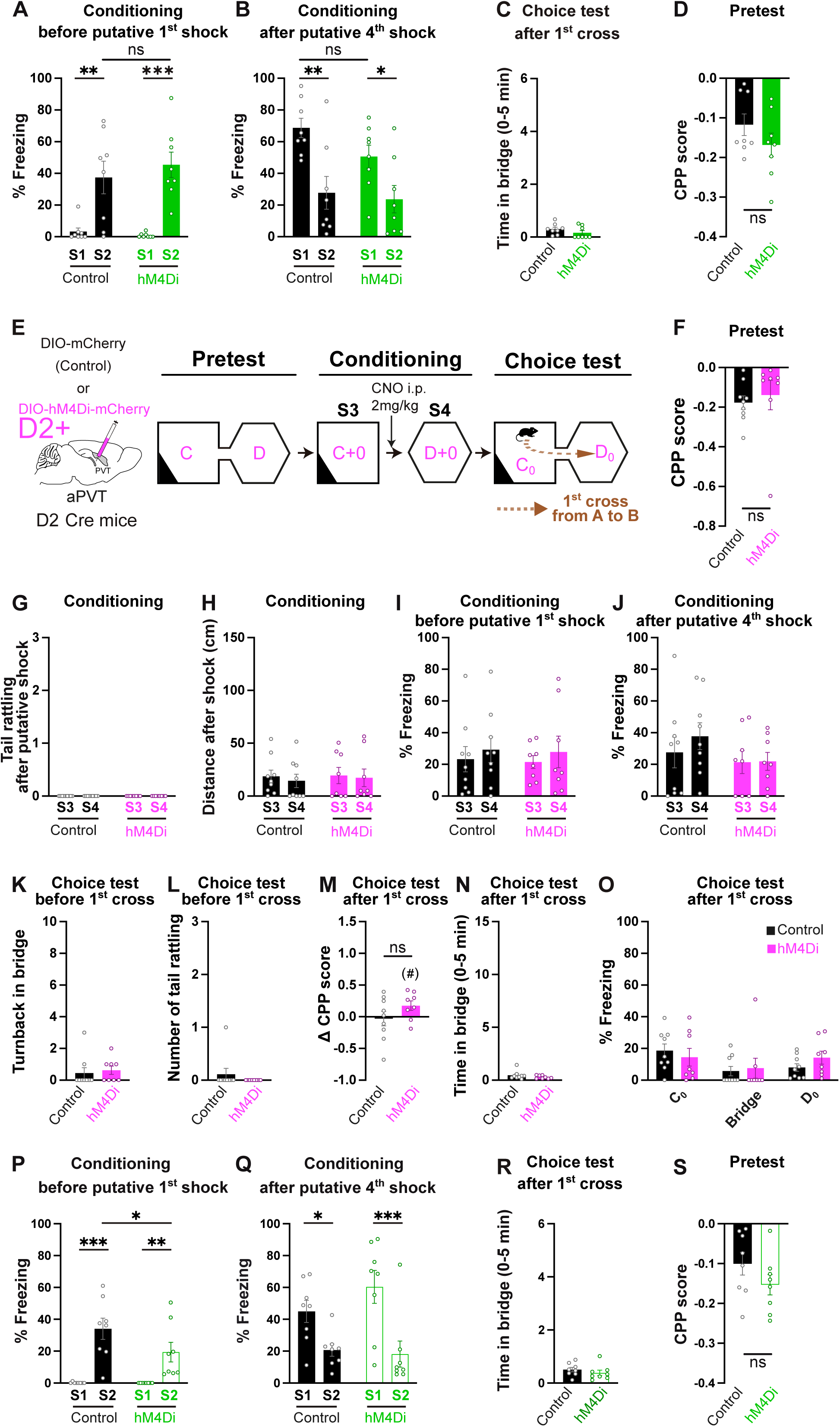

