## Supplementary material for "D2-expressing neurons of the anterior paraventricular nucleus of the thalamus as key players for modulating relative aversive and safety learning": Figure legends

**Figure 1. Mice adapt their defensive behavioral responses to safe and aversive conditioning.**

(A) Schema of the 1-day conditioned place preference (CPP) task for the control, safe and aversive groups. Contexts A (square) and B (circle) are connected via a bridge and contain electric shock grids. During conditioning session S1, Context A is paired with either 0 mA or four 0.6 mA electric shocks. During conditioning session S2 (30min after S1), Context B is paired with either 0 mA or four 0.3 mA electric shocks.

(B) Only mice from the aversive group increased their total freezing level during conditioning session S2 compared to S1.

(C) Safe and aversive groups, but not the control group, increased freezing before the 1$^{st}$ shock of the conditioning session S2 compared to S1.

(D) Freezing after the 4$^{th}$ shock significantly decreased for the safe group between conditioning sessions S1 and S2, while it was maintained for the aversive group.

(E) Only the aversive group exhibited tail rattling behavior immediately after shock delivery during conditioning session S2.

All data are mean ± SEM. Individual data points are displayed as dots. Control (n=11); safe (n=11); aversive (n=12); *p<0.05; **p<0.01; ***p<0.001; $^{##}$p<0.01 against 0, ns: non-significant.

See also Figure S1.

**Figure S1. Further analysis of defensive behavior during safe and aversive conditioning, related to Figure 1.**

(A) Picture of the conditioned place preference (CPP) setup for the control, safe and aversive groups. Contexts A (square) and B (circle) were connected via a bridge and contained electric shock grids.

(B) Mice from the control, safe and aversive groups did not freeze during pretest.

(C) Both safe and aversive groups increased their total freezing during the conditioning sessions S1 (left) and S2 (right) compared to control mice.

(D) While mice stayed a similar amount of time in the center of context A during conditioning session S1 (left), only mice from the aversive group spent significantly less time in the center during conditioning session S2 (right).

(E) Mean percentage of freezing ± SEM throughout S1 and S2 conditioning sessions for the control (A+0, B+0), safe (A+0.6, B+0) and aversive (A+0.6, B+0.3) groups. The four orange vertical lines indicate the timing of the four putative shocks delivered.

(F) Schema of the 1-day conditioned place preference (CPP) task for the control, safe and aversive groups indicating when shock events were delivered or their putative time of delivery during each 10-min conditioning session. The averaged percentage of freezing before 1$^{st}$ shock was calculated during min 1-3 while that after the 4$^{th}$ shock was calculated during min 9-10.

(G) No tail rattling was observed after the putative shocks delivered during the conditioning session S1 for the control, safe and aversive groups.

All data are mean ± SEM. Individual data points are displayed as dots. Control (n=11); safe (n=11); aversive (n=12); *p<0.05; **p<0.01; ***p<0.001.

**Figure 2. Mice use prior knowledge and display specific behavioral signatures to make appropriate safe and aversive choices.**

(A) During choice-test, mice from the control, safe and aversive groups are placed in their most preferred context (Context A, square). The 1st cross corresponds to the moment mice cross from Context A to Context B for the first time. A ΔCPP score is calculated from the 1st cross time event.

(B) Percentage of time mice spent in the bridge during choice-test before and after their 1st cross from context A to context B for the control, safe and aversive groups.

(C) Mice from the aversive group spent significantly more time in the bridge before their 1st cross from context $A_{0.6}$ to context $B_{0.3}$ than the control group.

(D) Mice from the aversive group made significantly more turnback in the bridge before making their 1st cross from context $A_{0.6}$ to context $B_{0.3}$ than the control and safe groups.

(E) Mice from the safe and aversive groups, but not from the control group, reverted their initial preference (measured during pretest) during the 15 min choice-test, by spending more time in the safest context B as indicated by a positive ΔCPP score (CPP score during the choice test minus CPP score during the pretest).

(F) Mice from the aversive group had a specific pattern of freezing during the entire choice-test compared to that of the safe and control groups in the context A (left), bridge (center) and context B (right). The aversive group froze more in the bridge than the safe and control groups. These mice also froze more in context B than the control group.

(G) Only mice from the aversive group spent more time in the bridge during choice test - pretest (Δ Time) after their 1st cross from context A to context B.

All data are mean ± SEM. Individual data points are displayed as dots. Control (n=11); safe (n=11); aversive (n=12); *$p<0.05$; **$p<0.01$; ***$p<0.001$. ## $p<0.01$ and ### $p<0.001$ against 0; ns: non-significant.

See also Figure S2.

**Figure S2. Further analysis of mice behavioral signatures during safe and aversive choices, related to Figure 2.**

(A) The first latency to cross from context A to context B during the choice test was similar between control, safe and aversive groups.

(B) The number of tail rattling before the 1st cross was similar between control, safe and aversive groups.

(C) Representative tracking path of mice from the control, safe and aversive groups during the entire choice test.

(D) CPP score during the pretest was similar between control, safe and aversive groups.

(E) The number of crossings between the two contexts during choice test was significantly lower for the safe and aversive groups compared to that of the control group.

(F) Mice from the control group (left panel) did not change their initial preference (measured during pretest) during choice test as indicated by a ΔCPP score (CPP score during the choice test minus CPP score during the pretest) around zero. Mice from the safe group (middle panel) reverted, especially during the first 5 minutes, their initial pretest preference during choice test as indicated by a positive

ΔCPP score. Mice from the aversive group (right panel) also changed their initial pretest preference during choice test, but with a positive ΔCPP score maintained throughout the entire 15 minutes.
(G) Only mice from the aversive group displayed a higher number of feces during the 15 min choice test.
All data are mean ± SEM. Individual data points are displayed as dots. For panel A to G: Control (n=11); safe (n=11); aversive (n=12); For panel H to J: Control (n=7); safe (n=9); relative aversive (n=9); absolute aversive (n=8); *p<0.05; ***p<0.001; * # p<0.05, ## p<0.01 and ### p<0.001 against 0. ns: non-significant.

**Figure 3. D2-expressing neurons (D2+) in the anterior PVT are preferentially recruited during both safe and aversive conditioning.**
(A) Schema of the 1-day conditioned place preference (CPP) task for the control home cage (HC), safe and aversive groups from D2 Cre-tagged mice (D2 Cre:Ai14 and D2 Cre ribotag mice). Mice were sacrificed 90 min after the conditioning session S2 to assess cFos expression in PVT brain sections.
(B) Schematics and representative image of the anterior PVT (aPVT, antero-posterior axis AP: -0.34) and posterior PVT (pPVT, antero-posterior axis AP: -1.34 showing examples of D2+ (in red) and cFos+ (in white) cells. Scale bar: 100 µm.
(C) Both safe and aversive conditioning led to a significant increase in cFos density (number of neurons per $mm^2$) in the aPVT compared to that of the pPVT.
(D) D2+ neurons in the aPVT were preferentially recruited after both safe and aversive conditioning compared to home cage control mice.
(E) D2+ and D2- neurons in the pPVT were similarly recruited by both safe and aversive conditioning compared to home cage control mice.
(F-G) The recruitment of D2+ (cFos+D2+ among D2+ cells), but not D2- (cFos+D2- among D2- cells), neurons during aversive conditioning was negatively correlated with the time spent in the bridge during the first 5 minutes of choice test after the 1$^{st}$ cross. Δ time in the bridge = time in bridge during choice test minus pretest for the first 5 min after the 1$^{st}$ cross from context A to context B for the aversive (up) and safe (down) groups. Line represents linear regression with confidence interval.
All data are mean ± SEM. Individual data points are displayed as dots. Home cage control (n=8); safe (n=10); aversive (n=10); *p<0.05; **p<0.01.
See also Figure S3.

**Figure S3. Further anatomical and behavioral analysis, related to Figure 3.**
(A) Safe and aversive groups increased freezing level before the 1$^{st}$ shock during conditioning session S2 compared to that of S1.
(B) Freezing after the 4$^{th}$ shock significantly decreased for the safe, but not the aversive, group between conditioning sessions S1 and S2.
(C) Representative cFos (white) immunostaining in the aPVT (up) and pPVT (down) of D2 Cre:Ai14 animals (D2+ neurons in red) in the home cage control, safe and aversive group. Scale bar=100 µm.

(D) A higher D2+ density (cell number per $mm^2$) was found in the pPVT than in the aPVT for the home cage control, safe and aversive groups.
(E) ΔCPP score (CPP score during choice test minus CPP score during the pretest) during the 15 min-choice test was similar between the safe and aversive groups.
(F) Mice in the aversive group spent more time than the safe group in the bridge during the first 5 minutes of choice test (normalized to the same time in the pretest). Δ time in the bridge = time in bridge during the choice test minus pretest for the first 5 minutes after the 1st cross from context A to context B.
All data are mean ± SEM. Individual data points are displayed as dots. Home cage control (n=8); safe (n=10); aversive (n=10); **p<0.01; ***p<0.001; (*) p=0.0584; (#) P=0.0511 and ### p<0.001 difference from 0; ns: non-significant.

**Figure 4. D2-expressing neurons are required for relative, but not absolute, aversive evaluation and choice.**
(A) D2 Cre mice were injected in the aPVT with either DIO-mCherry (n=8) or DIO-hM4Di-mCherry (n=8) viral vector (left panel). Representative image of mCherry expression in the aPVT of a D2 Cre mouse. Scale bar: 100 μm.
(B) Schema of the 1-day conditioned place preference (CPP) task for the aversive group. Control and hM4Di mice were i.p. injected with CNO (2mg/kg) 30 min before the conditioning session S2 in Context $B_{0.3}$ and were tested 1h later during a choice-test.
(C) hM4Di mice exhibited higher post-shock tail rattling during the conditioning session S2.
(D) The distance travelled immediately after shock decreased during conditioning session S2 compared to S1 for control mice. hM4Di mice did not show such reduced distance during S2.
(E) hM4Di mice showed increased turnbacks in the bridge before their 1st cross from context $A_{0.6}$ to context $B_{0.3}$.
(F) hM4Di mice exhibited increased tail rattling behavior before their 1st cross from context $A_{0.6}$ to context $B_{0.3}$.
(G) ΔCPP score (CPP score during choice test minus CPP score during pretest) was significantly decreased for hM4Di mice during the 15 min choice test, with hM4Di mice spending similar amount of time in the 2 contexts.
(H) hM4Di mice significantly spent more time in the bridge after their 1st cross from context $A_{0.6}$ to context $B_{0.3}$.
(I) hM4Di mice significantly decreased their freezing during the 15 min choice test in context $B_{0.3}$. All tests are comparisons between control and hM4Di for each compartment.
(J) A separated batch of D2 Cre mice was injected in the aPVT with either DIO-mCherry (n=8) or DIO-hM4Di-mCherry (n=7) viral vector.
(K) Schema of the contextual fear conditioning (CFC) task for the aversive group. Control and hM4Di mice were i.p. injected with CNO (2mg/kg) 30 min before the conditioning session S1 in Context $B_{0.3}$ and were tested 1h later during a retrieval test session.

(L) Control and hM4Di mice exhibited similar tail rattling after each shock of the conditioning session S1.
(M) Control and hM4Di mice similarly increased their freezing between the beginning (before 1st shock) versus the end (after 4th shock) of the conditioning session.
(N) Control and hM4Di mice exhibited similar tail rattling at the beginning (min1-3) of the retrieval test session.
(O) Total freezing percentage was similar between control and hM4Di mice during the retrieval test session.
All data are mean ± SEM. Individual data points are displayed as dots. Control for A-I (n=8), control for J-O (n=8), hM4Di for A-I (n=8), hM4Di for J-O (n=7); *$p<0.05$; **$p<0.01$; ***$p<0.001$; $^{###}$ $p<0.001$ difference from 0; ns: non-significant.
See also Figure S4.

**Figure S4. Further *ex vivo* and behavioral analysis and control experiments, related to Figure 4.**
(A) (Left) Example of whole-cell recording traces of control and hM4Di D2-expressing neurons of the aPVT expressing mCherry before and during 10μM CNO bath application. (Right) Bath application of 10μM CNO significantly decreased action potential frequency of aPVT D2+ neurons for the hM4Di-mCherry, but not for the control-mCherry. Mean action potential frequencies are compared between the same duration before and after CNO application. Control (n=6); hM4Di (n=5).
(B) Representative images depicting the injection site of the mCherry viral vector in the aPVT (AP: -0.34) and the projections of D2-expressing neurons in the Nucleus Accumbens (NAc, AP:0.86) and Amygdala (Amy, AP: -1.22), from left to right. Scale bar = 500 μm.
(C) CPP scores for hM4Di and control mice were similar during pretest.
(D) hM4Di mice froze similarly to control mice in S1 and S2 before the 1st shock.
(E) hM4Di mice froze similarly to control mice in S1 and S2 after the 4th shock.
(F) hM4Di and control mice had the same 1st latency to cross from context $A_{0.6}$ to context $B_{0.3}$.
(G) hM4Di and control mice spent the same amount of time in the bridge before their 1st cross from context $A_{0.6}$ to context $B_{0.3}$.
(H) A separate batch of D2 Cre mice was injected in the aPVT with either DIO-mCherry (n=9) or DIO-hM4Di-mCherry (n=9) viral vector.
(I) Schema of the 1-day conditioned place preference (CPP) task for the aversive group. Control and hM4Di mice were i.p. injected with CNO (2mg/kg) 30 min before the choice test session.
(J) ΔCPP score (CPP score during choice test minus CPP score during pretest) was similar between hM4Di and control mice during the 15 min choice test, with both groups preferring the $B_{0.3}$ context.
(K) Schema of the 1-day conditioned place preference task in which both A and B contexts were paired with 0.6mA electric shocks.
(L) Mice showed a similar number of tail rattling after shock delivery during conditioning session S1 and S2.
(M) The distance travelled immediately after shock was similar between conditioning S1 and S2.

(N) ΔCPP score (CPP score during choice test minus CPP score during pretest) during the 15 min choice test was not significant from zero.

(O) No difference of freezing in context $A_{0.6}$, bridge nor context $B_{0.6}$ was found during the choice test.

(P) A separate batch of D2 Cre mice was injected in the aPVT with either DIO-mCherry (n=8) or DIO-hM4Di-mCherry (n=7) viral vector.

(Q) Schema of the 1-day conditioned place preference task in which no shock was delivered in Context C+0mA and 4 shocks of 0.3mA were delivered in Context D+03mA. The same mice first went through contextual fear conditioning (session 1) in context $B_{0.3}$ one week before (hence the sessions S2 and S3). Control and hM4Di mice were i.p. injected with CNO (2mg/kg) 30 min before the conditioning session S3.

(R) hM4Di and control mice exhibited higher tail rattling after putative shock during conditioning session S4 compared to S3.

(S) No change in tail rattling behavior was observed during choice test in the hM4Di and control mice before their 1st cross from context $D_{0.3}$ to context $C_0$.

(T) ΔCPP score (CPP score during choice test minus CPP score during pretest) during the 15 min choice test was negative for both groups, indicating they both successfully avoided $D_{0.3}$.

All data are mean ± SEM. Individual data points are displayed as dots. Control for C-G (n=8), for H-J (n=9), for P-T (n=8); hM4Di for C-G (n=8), for H-J (n=9), for P-T (n=7); control mice for K-O (n=8); **p<0.01; ***p<0.001; # p<0.05 and ## p<0.01 against 0. ns: non-significant.

**Figure 5. aPVT D2- (minus) neurons of the aPVT are not required for aversive evaluation and choice.**

(A) D2 Cre mice were injected in the aPVT with either FLEx-OFF-mCitrine (n=8) or FLEx-OFF-hM4Di-mCitrine (n=9) viral vector (left panel). Representative image of mCitrine viral expression in the aPVT in a D2 Cre mouse. Scale bar: 100 µm.

(B) Schema of the 1-day conditioned place preference (CPP) task for the aversive group. Control and hM4Di mice were i.p. injected with CNO (2mg/kg) 30 min before the conditioning session S2 in Context $B_{0.3}$ and were tested 1h later during a choice-test.

(C) hM4Di and control mice exhibited similar tail rattling after each shock of the conditioning sessions S1 and S2.

(D) The distance travelled immediately after shock decreased during conditioning S2 compared to S1 for both hM4Di and control mice.

(E) Similar number of turnback in the bridge zone and (F) tail rattling during choice test session before the 1st cross from context $A_{0.6}$ to context $B_{0.3}$ for hM4Di and control mice.

(G) Similar ΔCPP score (CPP score during the choice test - CPP score during the pretest) during the 15 min choice test between hM4Di and control mice, with both groups spending more time in context $B_{0.3}$.

(H) No change in time in the bridge after the 1st cross from context $A_{0.6}$ to context $B_{0.3}$ during the choice test session between hM4Di and control mice.

(I) No change in freezing level in none of the compartments ($A_{0.6}$, bridge and $B_{0.3}$) between hM4Di and control mice.

All data are mean ± SEM. Individual data points are displayed as dots. control (n=8) and hM4Di (n=9). **p<0.01; ***p<0.001; ## p<0.01 and ### p<0.001 against 0. ns: non-significant.

See also Figure S5.

**Figure S5. Further *ex vivo*, anatomical and behavioral analysis, related to Figure 5.**

(A) D2 Cre mice were injected in the aPVT with either FLEx-OFF-mCherry (n=2) or FLEx-OFF-hM4Di-mCherry (n=2) viral vector.

(B) Example of whole-cell recording traces of control and hM4Di D2- (minus) neurons of the aPVT expressing mCherry before and during 10μM CNO bath application. Bath application of 10μM CNO significantly decreased action potential frequencies of aPVT D2-neurons for the hM4Di-mCherry, but not for the control-mCherry. Mean action potential frequency are compared between the same duration before and after CNO application. Control (n=6); hM4Di (n=6).

(C) Representative image of mCitrine (green, D2-) viral expression in the aPVT of a D2 Cre:Ai14 mouse. D2+ cells express tdTomato (magenta, D2+). Scale bar: 100 μm.

(D) 3 D2 Cre:Ai14 mice were injected with either FLEx-OFF-mCitrine (n=1) or FLEx-OFF-hM4Di-mCitrine (n=2) viral vector. We found low overlap (3±0.38%) between D2-expressing (D2+) and D2 minus (D2-) neurons (109 cells out of 3475 cells were mCitrine+ (green, D2-) and tdTomato+ (magenta, D2+).

(E) Both hM4Di and control mice increased their freezing before the 1$^{st}$ putative shock from S1 to S2. Freezing in S2 was lower in hM4Di mice than in control mice.

(F) hM4Di and control mice increased their freezing after the 4$^{th}$ putative shock from S1 to S2.

(G) hM4Di and control mice showed the same 1$^{st}$ latency to cross from context $A_{0.6}$ to context $B_{0.3}$.

(H) hM4Di and control mice spent the same amount of time in the bridge before their 1$^{st}$ cross from context $A_{0.6}$ to context $B_{0.3}$.

All data are mean ± SEM. Individual data points are displayed as dots. control for A-B (n=2), control for C-D (n=1), control for E-I (n=8); hM4Di for A-B (n=2), hM4Di for C-D (n=2), hM4Di for E-I (n=9); *p<0.05; **p<0.01; ***p<0.001; ns: non-significant.

**Figure 6. Inactivating aPVT D2+ neurons increases contextual aversiveness and related threat-predicting behavioral traits.**

(A) Schema of the 1-day conditioned place preference task in which both A and B contexts were both paired with 0.3mA. Control and hM4Di mice were i.p. injected with CNO (2mg/kg) 30 min before the conditioning session S2.

(B) hM4Di mice displayed a greater number of tail rattling during conditioning session 2 compared to control mice.

(C) Freezing after the 4$^{th}$ shock in S2 was similar between hM4Di and control mice.

(D) hM4Di displayed more turnback in the bridge before the 1$^{st}$ cross from context $A_{0.3}$ to context $B_{0.3}$ than control mice.

(E) hM4Di and control mice showed a similar 1st latency to cross from context $A_{0.3}$ to context $B_{0.3}$.

(F) hM4Di mice and control mice spent a similar amount of time in the bridge before their 1st cross between context $A_{0.3}$ to context $B_{0.3}$.

(G) hM4Di and control mice froze similarly in context $A_{0.3}$, bridge and $B_{0.3}$ during choice test. All tests are comparisons between control and hM4Di for each compartment.

(H) ΔCPP score (CPP score during choice test - CPP score during pretest) during the 15 min choice test is significantly lower and not different from 0 in hM4Di mice compared to that of control mice.

All data are mean ± SEM. Individual data points are displayed as dots. control (n=8); hM4Di (n=7); (*) p=0.0565; *p<0.05; ***p<0.001; $^{\#}$p<0.05 against zero; ns: non-significant.

See also Figure S6.

**Figure S6. Further behavioral analysis and control experiments, related to Figure 6.**

(A) hM4Di and control mice traveled the same distance after shock between S1 and S2.

(B) Increased freezing from S1 to S2 before the 1st shock was similar between hM4Di and control mice.

(C) Tail rattling before the 1st cross from context $A_{0.3}$ to context $B_{0.3}$ was similar between hM4Di and control mice.

(D) Time mice spent in the bridge after their 1st cross from context $A_{0.3}$ to context $B_{0.3}$ was similar between hM4Di and control mice.

(E) CPP scores for hM4Di and control mice were similar during pretest.

All data are mean ± SEM. Individual data points are displayed as dots. control (n=8); hM4Di (n=7); ***p<0.001. ns: non-significant.

**Figure 7. Inhibiting aPVT D2+, but not D2- neurons, increases contextual safeness and related safety-predicting behavioral traits.**

(A, J) Schema of the 1-day conditioned place preference task for the safe groups. (A) Control (AAV-DIO-mCherry) and hM4Di mice (AAV-DIO-hM4Di-mCherry) and (J) Control (AAV-FLEx-OFF-mCitrine) and hM4Di mice (AAV-FLEx-OFF-hM4Di-mCitrine) were i.p. injected with CNO (2mg/kg) 30 min before the conditioning session S2.

(B, K) hM4Di mice did not change their number of tail rattling during conditioning S1 and S2.

(C, L) hM4Di and control mice similarly showed a decreased distance after putative shock time event between S1 and S2.

(D, M) hM4Di and control mice showed a similar number of turnback in the bridge before 1st cross from context $A_{0.6}$ to context $B_0$.

(E, N) hM4Di and control mice exhibited a similar number of tail rattling before 1st cross from context $A_{0.6}$ to context $B_0$.

(F, O) hM4Di mice did not change their 1st latency to cross from context $A_{0.6}$ to context $B_0$.

(G) Inactivating aPVT D2+, (P) but not D2- neurons, significantly increased their 2nd latency to cross back from the context $B_0$ to the context $A_{0.6}$ during choice test.

(H) During choice test, inactivating aPVT D2+, (Q) but not D2- neurons, significantly decreased freezing in context $A_{0.6}$, it increased freezing in context $B_0$, with no change in freezing in the bridge. All tests are comparisons between control and hM4Di for each compartment.

(I) Inactivating aPVT D2+, (R) but not D2-, neurons significantly increased ΔCPP score (CPP score during choice test minus CPP score during pretest) during the 15 min choice test.

All data are mean ± SEM. Individual data points are displayed as dots. For D2+ and D2- experiments: Control (n=8); hM4Di (n=8); *p<0.05; **p<0.01; $^{##}$p<0.01; $^{###}$p<0.001; ns: non-significant.

See also Figure S7.

**Figure S7. Further behavioral analysis and control experiments, related to Figure 7.**

(A) hM4Di and control mice showed a similar increase of freezing from S1 to S2 before the 1$^{st}$ putative shock.

(B) hM4Di and control mice showed a decreased freezing between S1 and S2 after the 4$^{th}$ putative shock.

(C) hM4Di mice did not change the time spent in the bridge after their 1$^{st}$ cross from context $A_{0.6}$ to context $B_0$.

(D) CPP scores for hM4Di and control mice were similar during pretest.

(E) Schema of the 1-day conditioned place preference task in control condition in which none of the contexts (C and D) were paired with shocks. Note that mice went through the $A_{0.3}$-$B_{0.3}$ conditioning paradigm and performed the 0-0 control task one week later (hence the sessions S3 and S4). Control (n=9) and hM4Di (n=8) mice were i.p. injected with CNO (2mg/kg) 30 min before the conditioning session S4.

(F) CPP scores for hM4Di and control mice were similar during pretest.

(G) hM4Di mice did not exhibit tail rattling after putative shock time events during S3 and S4.

(H) hM4Di and control mice showed a similar distance after putative shock time event between S3 and S4.

(I) hM4Di mice did not change their freezing level between S3 and S4 before the 1$^{st}$ putative shock time event.

(J) hM4Di mice did not change their freezing level between S3 and S4 after the 4$^{th}$ putative shock time event.

(K) hM4Di and control mice displayed similar number of turnbacks in the bridge before the 1$^{st}$ cross from context $C_0$ to context $D_0$.

(L) hM4Di and control mice exhibited a similar number of tail rattling before 1$^{st}$ cross from context $C_0$ to context $D_0$.

(M) ΔCPP score (CPP score during choice test minus CPP score during pretest) was similar between hM4Di and control mice during the 15 min choice test.

(N) hM4Di mice did not change the time spent in the bridge after their 1$^{st}$ cross between context $C_0$ to context $D_0$.

(O) hM4Di and control mice did not change their freezing level in context $C_0$, bridge nor context $D_0$ during choice test. All tests are comparisons between control and hM4Di for each compartment.

(P) Both hM4Di and control mice increased their freezing before the 1st putative shock from S1 to S2. Freezing in S2 was lower in hM4Di mice than in control mice.

(Q) hM4Di and control mice decreased their freezing after the 4th putative shock from S1 to S2.

(R) hM4Di and control mice spent the same amount of time in the bridge after their 1st cross.

(S) CPP scores for hM4Di and control mice were similar during pretest.

All data are mean ± SEM. Individual data points are displayed as dots. Control (n=8) for A-D; Control (n=9) for G-O; Control (n=8) for P-S; hM4Di (n=8) for A-D; hM4Di (n=8) for G-O; hM4Di (n=8) for P-S; *p<0.05; **p<0.01; ***p<0.001; (#) P=0.0543 against 0; ns: non-significant.
